# Single-nucleus multiome ATAC and RNA sequencing reveals the molecular basis of thermal acclimation in *Drosophila melanogaster* embryos

**DOI:** 10.1101/2025.01.08.631745

**Authors:** Thomas S. O’Leary, Emily E. Mikucki, Sumaetee Tangwancharoen, Joseph R. Boyd, Sara Helms Cahan, Seth Frietze, Brent L. Lockwood

## Abstract

Embryogenesis is remarkably robust to temperature variability, yet the homeostatic mechanisms that offset thermal effects during early development remain poorly understood. We measured how acclimation shifts the upper thermal limit and profiled chromatin accessibility and gene expression in *D. melanogaster* embryos using single-nucleus multiome sequencing. Acclimation preserved shared primordial cell types yet rapidly shifted heat tolerance. Cool-acclimated embryos showed a homeostatic response characterized by increased accessibility at binding motifs for the transcriptional activator *Zelda*, along with enhanced activity of gene regulatory networks in a subset of primordial cell types including the gut, muscle, and nervous system. Cool-acclimated embryos also had higher expression of ribosomal and oxidative phosphorylation genes, reflecting a coordinated increase in translation and energy production to buffer slower biochemical kinetics in the cold. Our results indicate that maintaining developmental robustness across temperature requires homeostatic gene expression responses that may carry metabolic costs constraining upper thermal limits.

## Introduction

Early embryonic development in *Drosophila* involves the spatially and temporally coordinated activation of zygotic transcription, producing precise gene expression patterns across the embryo (*1–4*). These morphogenetic gradients drive key transitions, such as gastrulation and the primordial specification of tissues, that result in consistent cell-type compositions despite genetic and environmental variation (*4–13*), a phenomenon known as developmental robustness. However despite this well-established paradigm (*14*), and decades of research on embryonic morphology and gene regulatory networks (*15*), surprisingly little is known about how early embryos physiologically respond to environmental variability (*16*).

This gap is critical because developmental systems evolved under variable natural conditions, not the controlled environment of the lab. In nature, embryos experience fluctuating temperatures and their physiological responses can influence species survival (*17–22*). Acclimation of upper thermal limits in particular, may shape a species’ vulnerability to climate change (*23*). Because embryos cannot behaviorally thermoregulate and have a lower thermal limit than later life stages (*5*, *17*), even modest shifts in thermal tolerance at this stage could significantly affect species persistence (*17*, *24*).

Early embryonic development involves three key complex cellular and molecular events:

(i) the transition from maternal to zygotic control of gene expression (*3*, *25*, *26*), (ii) cellularization, gastrulation, and cell migration (*10*, *13*, *27*), and (iii) cell type specification and tissue differentiation (*28*, *29*). Although these processes depend on complex, temperature-sensitive interactions between proteins, DNA, and other macromolecules, they proceed normally across a range of temperatures (*7*, *30–33*).

The ability to maintain consistent morphology and cell-type composition despite temperature variation is remarkable, given that temperature affects the kinetics of all biochemical reactions, including transcription and translation (*34–37*). Because thermal effects on biological rates are so pervasive, organisms rely on homeostatic mechanisms that actively regulate gene or protein expression (*38–40*) to maintain performance across temperatures (*32*, *41*, *42*). While thermal acclimation responses are well documented in later developmental stages and adulthood (*43–46*), it remains unclear whether similar homeostatic mechanisms operate during embryogenesis. This knowledge gap is likely due to the predominance of studies conducted under static laboratory conditions (*28*, *47*, *48*).

Here, we measured heat tolerance in *D. melanogaster* embryos reared to Bownes’ stage 11 across a 12°C acclimation range to assess how thermal acclimation influences survival under extreme heat. We then used single-nucleus multiome sequencing (ATAC-seq and RNA-seq) to profile chromatin accessibility and gene expression at cell-type resolution, allowing us to characterize both cellular composition and molecular responses to temperature during embryogenesis. We selected stage 11 as a critical developmental window when embryos transition from multipotency to terminal cell fate specification (*48*), providing an opportunity to examine mechanisms that preserve developmental progression despite thermal variation. Given the strong connection between temperature and ectotherm physiology, we hypothesized that embryonic thermal acclimation would affect (i) heat tolerance and (ii) cellular and developmental homeostatic mechanisms. Our findings reveal previously undescribed physiological responses during early development and highlight the ecological relevance of thermal plasticity in shaping developmental robustness.

## Results

### Thermal acclimation rapidly alters embryonic heat tolerance

To assess the phenotypic effects of thermal acclimation, we reared embryos for 9 hours at 18°C, 5 hours at 25°C, or 3 hours at 30°C (see Methods), then measured hatching success following a 45-minute heat shock at 38.75°C, a temperature near the upper thermal limit of embryos at this stage (data not shown). Acclimation temperature had a significant effect on heat tolerance (**Fig. 1**; quasi-binomial logistic regression, p < 1 × 10^-5^), with survival increasing at higher acclimation temperatures. Only 21.0% of the 18°C-acclimated embryos hatched, compared to 42.6% and 49.5% for the 25°C and 30°C-acclimated groups, respectively. Post hoc tests confirmed that the 18°C-acclimated embryos were significantly more heat sensitive than both the 25°C and 30°C-acclimated embryos (Tukey-adjusted pairwise Wald test on estimated marginal means; p < 1 × 10^-4^ for both), while the 25°C and 30°C groups did not differ significantly (p = 0.18).

**Fig. 1.**
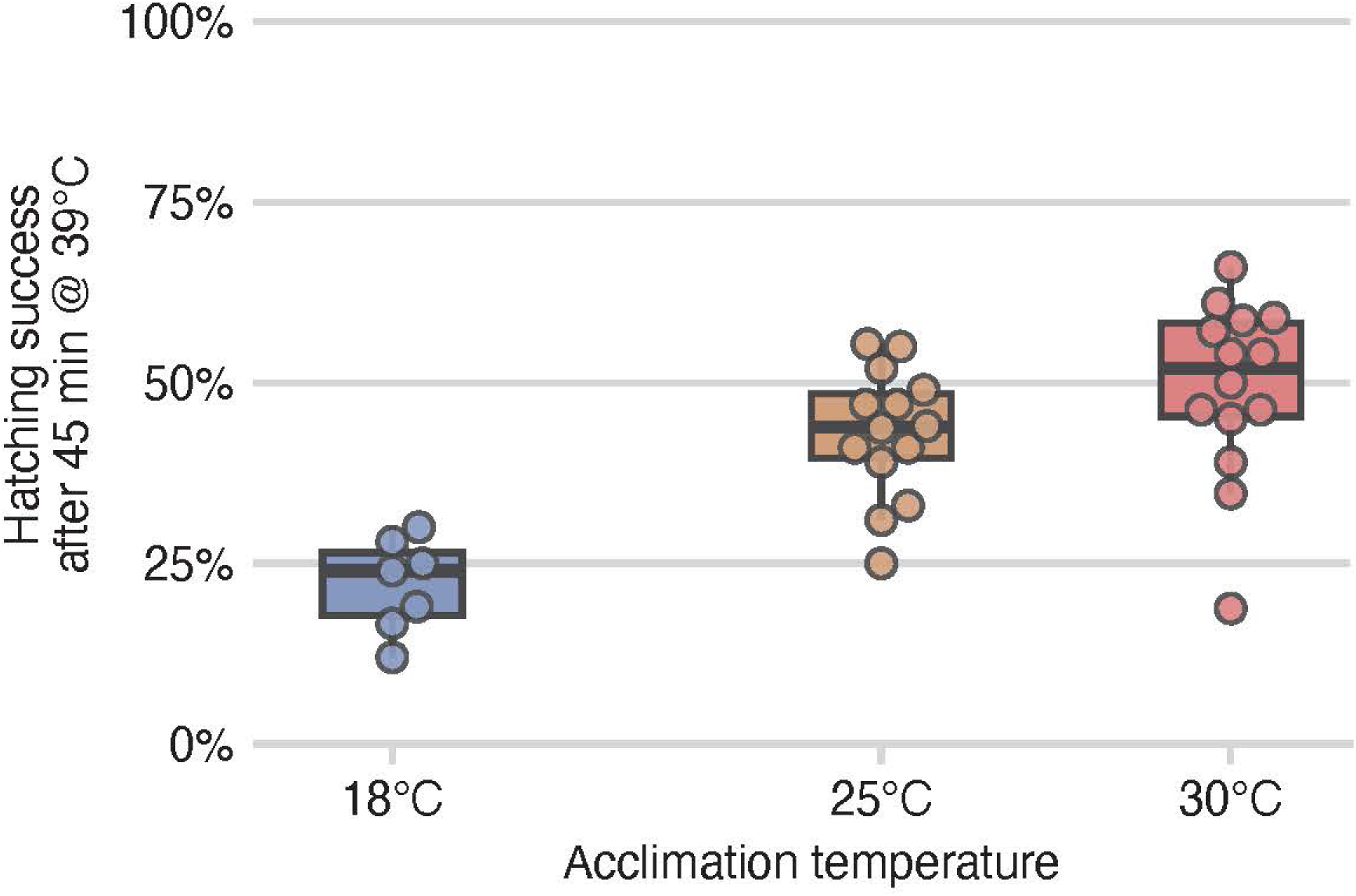
Thermal acclimation of early embryos led to differential acute heat shock tolerance. The hatching success of embryos following an acute heat shock of 38.75°C for 45 minutes. Hatching success increased with increasing acclimation temperature (quasi-binomial logistic regression; p < 1 × 10^-5^). Individual dots represent replicate grids of approximately 50 to 100 embryos, total number of eggs per acclimation temperature: 520, 1272, 1145 for 18°C, 25°C, and 30°C respectively.

### Multiome profiling of thermal acclimation in embryos

We focused our multiomics analysis on the 18°C and 25°C acclimation conditions, which showed the greatest difference in heat tolerance despite only a 7°C shift in temperature.

For each condition we pooled 50 Bownes’ stage 11 embryos per replicate (two replicates per condition) targeting approximately 4,000 high-quality nuclei per sample for the 10x Genomics single-nucleus ATAC and RNA sequencing. Sequencing yielded 262 million ATAC-seq read pairs and 214 million RNA-seq reads, mapping to 464,722 barcodes with reads in both modalities (**Fig. S1C**). After filtering for low read counts, high mitochondrial or ribosomal content, and predicted multiplets, we retained a total of 10,390 high-quality nuclei (5,298 from the 18°C and 5,092 from the 25°C; **Fig. S1, A and B**). Per nucleus, there was a median of 983 RNA-seq reads and 5,688 ATAC-seq reads (**Fig. S2, A and B**). Across the dataset, we detected 11,271 peaks and 9,007 genes, with a median of 2,206 peaks and 586 genes per nucleus (**Fig. S2, C and D**). Differential accessibility and expression were conducted using MAST (*49*), which accounts for the detection rate at the nucleus level. This allowed testing of 5,017 to 6,782 peaks (median of 6,100; **Table S1**) and 1,217 to 2,004 genes (median of 1,598; **Table S2**) per cell type.

### Cell-type composition is conserved across acclimation temperatures

After quality control, the remaining 10,390 nuclei were visualized using Uniform Manifold Approximation and Projection (UMAP), integrating both ATAC and RNA data via a weighted k-nearest neighbor approach. Nuclei from the 18°C and 25°C acclimation treatments showed substantial overlap, indicating conserved developmental trajectories despite temperature differences (**Fig. 2A**). We identified 12 distinct clusters using a shared nearest neighbor algorithm applied to the integrated ATAC and RNA data (**Fig. 2B**). To annotate these clusters, we identified 1,564 cluster-specific marker genes from the expression data (**Fig. 2E; Fig. S4**). Following the approach of Calderon *et al.*, (2022), we assigned biological labels to clusters using a Fisher’s enrichment test for overlap with annotations from the Berkeley Drosophila Genome Project RNA *in situ* hybridization database (*50*). Final annotations were made manually, guided by these enrichment results (**Fig. 2C**).

**Fig. 2.**
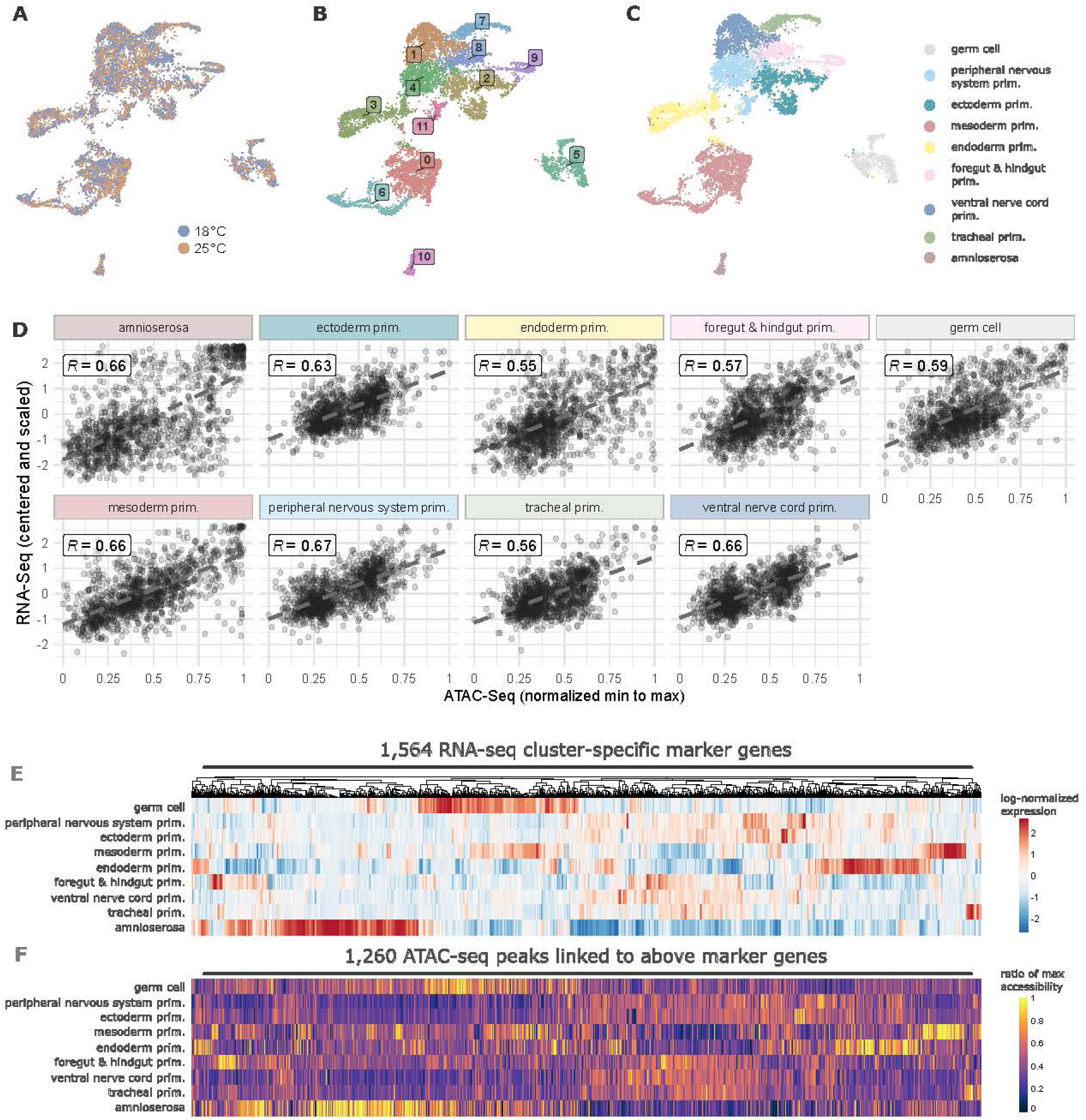
Nuclei cluster similarly across acclimation states and were annotated to cell-type labels using identified marker genes, with a strong concordance between ATAC-seq and RNA-seq data. (A-C) A UMAP representation of the single-nucleus multiome data. Each dot represents an individual nucleus characterized by integrative profiling of gene expression (RNA) and chromatin accessibility (ATAC) condensed into a two-dimensional representation. UMAPs are colored by three different representations, acclimation treatment (**A**), cluster number (**B**), and annotated cell types (**C**). (**D**) Each point on the scatter plot represents a marker gene and the top linked peak identified in the ATAC-seq data. The ATAC-seq data was normalized using minimum to maximum normalization (*i.e.,* min-max scaling) and the RNA-seq data was centered and scaled (*i.e.,* z-score). Each panel represents different primordial cell types. Heatmaps of the same data with RNA-seq cluster specific marker genes (**E**) and the top linked peak in ATAC-seq data (**F**).

### Chromatin accessibility corroborates RNA-based cell-type annotations

To validate RNA-based cell-type annotations, we assessed the concordance between chromatin accessibility (ATAC-seq) and gene expression profiles across clusters. The multiome data revealed strong agreement between RNA-defined marker genes and corresponding chromatin accessibility patterns. Of the 1,564 cluster-specific marker genes, 1,260 (80%) had nearby ATAC-seq peaks significantly correlated with expression. The top linked peaks (lowest adjusted p-value) showed clear cell-type-specific accessibility matching gene expression patterns (**Fig. 2F**). This concordance was consistent across all marker genes and their linked accessibility peaks, with cell-type-specific correlations ranging from 0.55 and 0.66 (**Fig. 2D**). To illustrate this in detail, we visualized ATAC-seq coverage tracks for three representative marker genes, *btl*, *Mef2*, and *otp*, highlighting matched accessibility and expression in tracheal primordium, mesoderm primordium, and foregut & hindgut primordium, respectively (**Fig. S5**).

### Zelda and HSF motifs show differential accessibility between acclimation temperatures

To assess broad changes in chromatin accessibility across conditions, we performed pseudobulk analysis by combining all nuclei and identifying differentially accessible regions (DARs) in the ATAC-seq data. This analysis identified 142 DARs between acclimation temperatures, with 84 regions (59%) with increased accessibility in the cool-acclimated embryos and 58 (41%) more accessible in the 25°C-acclimated embryos (**Fig. S6A**). We next conducted motif enrichment analysis on these DARs to identify overrepresented TF binding motifs. Out of the 150 motifs tested, only one target motif corresponding to the pioneer transcription factor ZLD (*Zelda*) essential for early zygotic genome activation was significantly enriched. Among the 710 ZLD motif-containing peaks detected in at least 10% of nuclei, 33 were differentially accessible between treatments, with most showing increased accessibility in the cool-acclimated embryos (**Fig. 3A**). GO enrichment of these 33 ZLD-target DARs revealed developmental processes such as axis elongation and cell migration (**Fig. S6C**). We highlight three ZLD-target DARs linked to genes with known developmental roles. *smash* (**Fig. 3B**) encodes a microtubule-and junction-associated protein highly expressed in late embryogenesis and pupation (*51*, *52*). *halo* (**Fig. 3C**) encodes a kinesin co-factor involved in histone variant exchange during early embryogenesis (*53*), with potential implications for chromatin remodeling. *Nrt* (**Fig. 3D**) encodes Neurotactin, a cell adhesion protein required for gastrulation and ventral furrow formation (*54*, *55*).

**Fig. 3.**
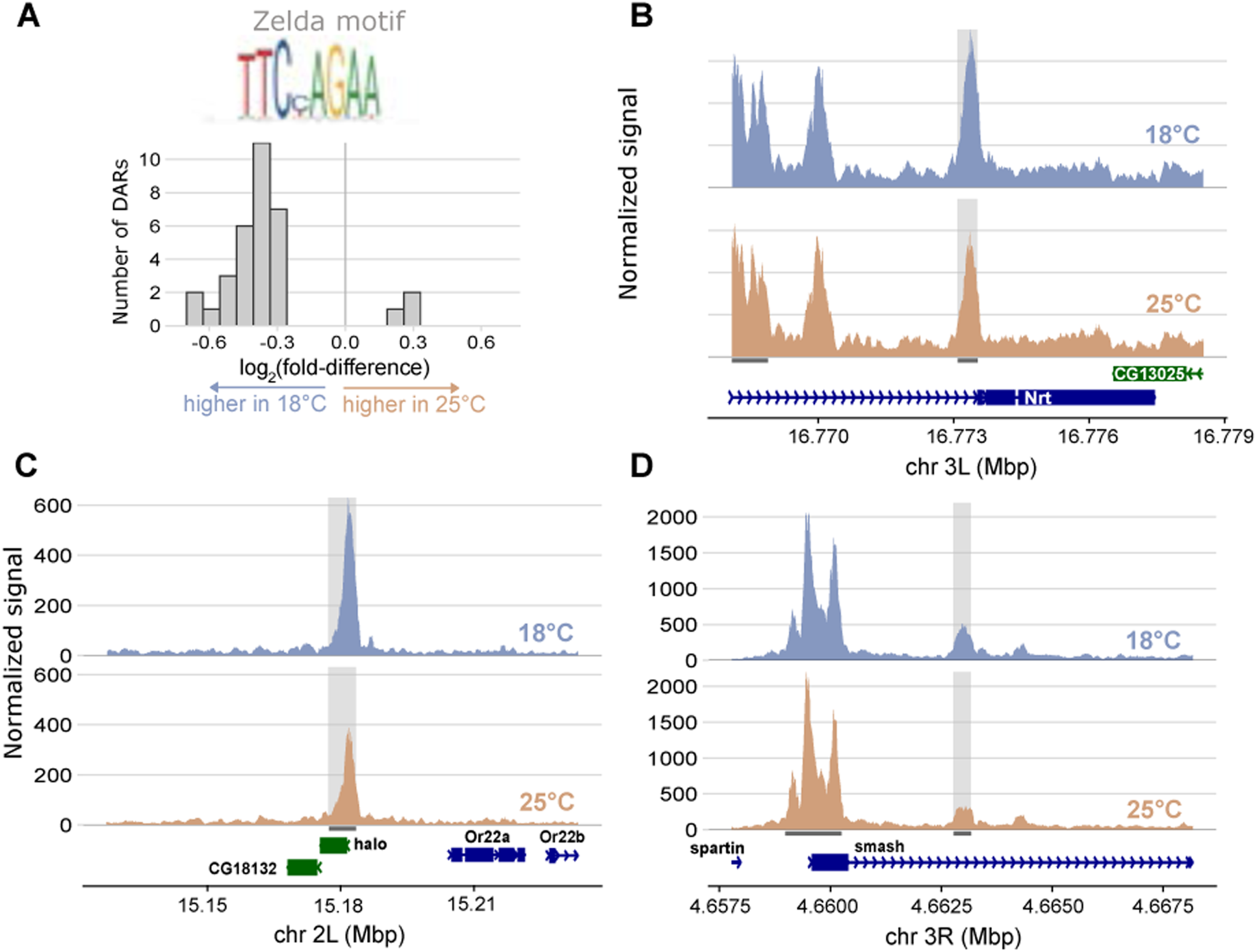
The majority of pseudobulk DARs with *Zelda*-target motifs show increased accessibility in the 18°C cool-acclimated embryos. (**A**) Histogram of log2(fold-difference) between acclimation treatments of the 33 pseudobulk DARs that contain the Zelda-target motif. Pseudobulk ATAC-seq coverage plots for three representative Zelda-target motif containing DARs (**B-D**). Within each panel containing the coverage plots, the top trace, colored blue, represents the accessibility of the 18°C-acclimated embryos, and the bottom trace, in orange, represents the 25°C-acclimated embryos. Grey shading indicates the peak regions with differential accessibility between acclimations.

Because chromatin accessibility is known to be established prior to heat shock (*56*), we also examined accessibility at peaks containing heat shock factor (HSF) motifs. HSF is the transcription factor that activates molecular chaperones, or heat shock proteins (HSP), in response to thermal stress (*57*). Of the 518 HSF-target peaks detected in at least 10% of nuclei, 12 were differentially accessible between acclimation conditions (**Fig. 3F**).

However, none of these DARs were linked to canonical HSP genes in our dataset (*Hsp23, Hsp26, Hsp27, Hsp60A, Hsp67Ba, Hsp70Ab,* and *Hsp83*). Overall, HSF-target motifs were less accessible in 25°C-acclimated embryos (**Fig. S7A**). We visualized two representative HSF-motif-containing DARs (**Fig. S7, B and C**), both associated with genes involved in transcriptional activation during key developmental windows. *pre-lola-G* is highly expressed at the developmental stage examined and is known to trans-splice with isoforms of *lola*, a gene that promotes chromatin accessibility of target promoters later in embryogenesis (*58*). *elB* encodes a transcription factor critical to tracheal formation, with peak expression late in embryogenesis (*59*, *60*).

### Gene regulatory networks reveal differential responses to thermal acclimation

Our analysis revealed that gene regulatory networks remained stable across acclimation temperature, with consistent transcription factor expression and motif accessibility across cell types (**Fig. 4A**). However, a subset of GRNs showed a clear acclimation-dependent changes (**Fig. 4B**). Using the SCENIC+ pipeline (*61*), we identified 41 enhancer-driven GRNs (referred to as eRegulons in SCENIC+ pipeline; in this manuscript we use the term GRN). This GRN-based approach enabled us to identify transcription factors, motifs, and downstream targets that define cell-type-specific regulatory programs during development, and to test whether these networks were modulated by thermal acclimation. Of the 41 GRNs with strong transcription factor-cistrome correlation (|rho| > 0.5; **Fig. S8**), 29 acted as activators (motif opening and target gene upregulation), and 12 as repressors (motif closing and target gene downregulation) (**Fig. 4A**). While most networks were unaffected by temperature, we identified 18 transcription factors that were differentially expressed between acclimation treatments within specific cell types. Of these, 10 were part of GRNs that were differentially regulated (hereafter diffGRNs), defined by coordinated changes in motif accessibility and target gene expression (**Fig. 4B**). For example, *Blimp-1*, a transcription factor expressed in the foregut and hindgut primordium, was upregulated in 18°C-acclimated embryos (log_2_ fold-difference = 1.41). This was accompanied by increased accessibility of its target motifs (one-sample t-test, p = 5.4 × 10^-13^) and increased expression of its target genes (p = 1.3 × 10^-9^). Among the ten diffGRNs, four were found in the mesoderm primordium, three in the peripheral nervous system primordium, two in the ventral nerve cord primordium, and one in the foregut and hindgut primordium. Of the eight diffGRNs acting as activators, six showed increased target gene expression in the cool-acclimated embryos.

**Fig. 4.**
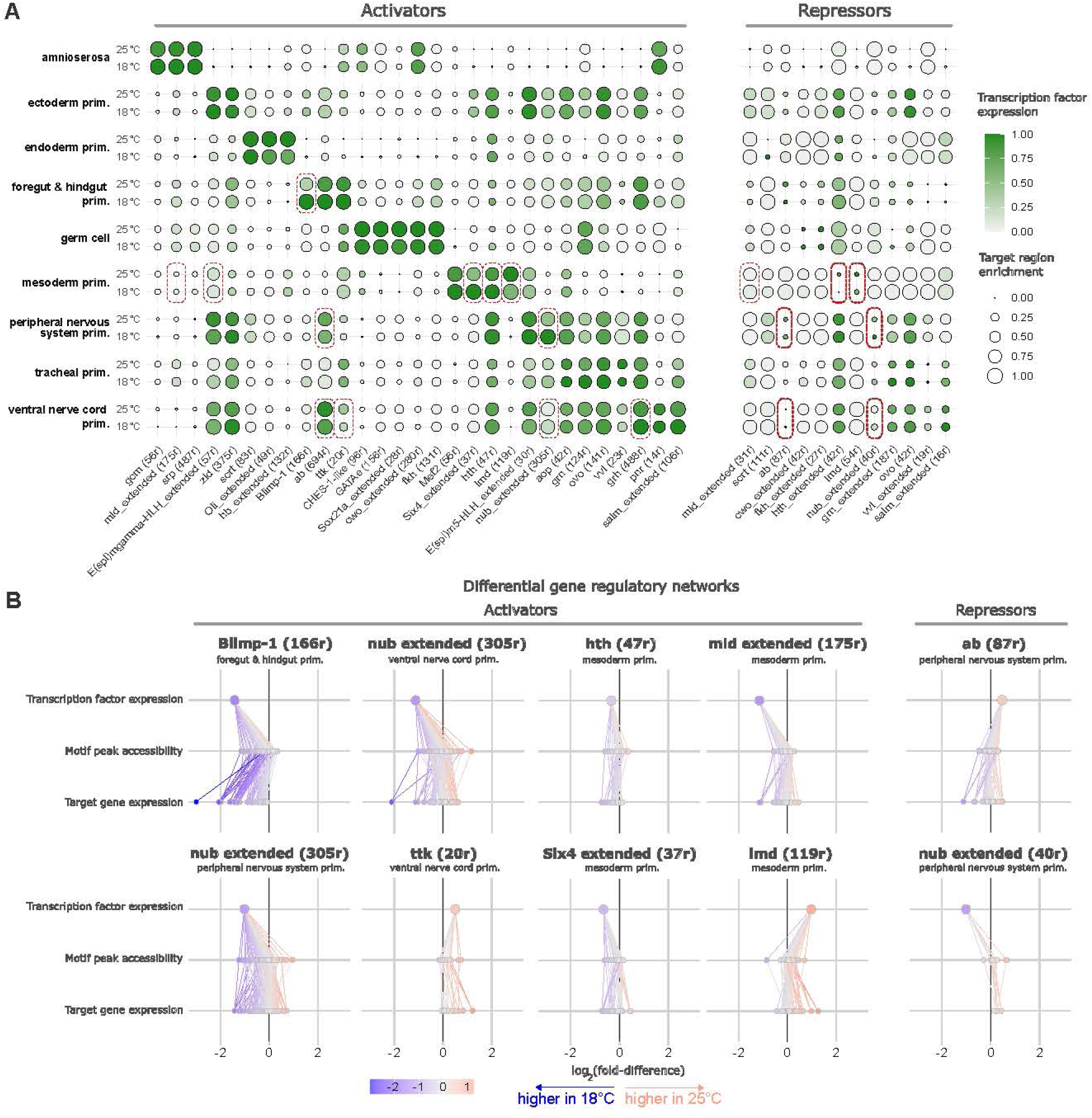
Ten cell-type-specific GRNs were altered with acclimation temperature, while the rest remained stable despite variation in environmental temperature. (**A**) SCENIC+ identified GRNs separated out by cell-type and acclimation, 29 activator GRNs are on the left panel and 12 repressors are on the right panel. The green color represents normalized transcription factor expression and area of the dot represents target region enrichment with greater size indicating more accessible chromatin. Differentially expressed transcription factors are indicated with red boxes. (**B**) Ten differential gene regulatory networks (diffGRNs) plotted with log_2_ fold-difference for each node in the network. Colored by log_2_ fold-difference with red indicating higher expression or accessibility in the 25°C-acclimated embryos and blue indicating higher in the 18°C cool-acclimated embryos. diffGRNs labeled as activators are on the left, and repressors are on the right.

### WGCNA reveals acclimation response of co-expressed genes

To identify global gene expression patterns and temperature-dependent transcriptional responses, we performed high-dimensional weighted gene co-expression network analysis (hdWGCNA) on the single-nucleus RNA-seq data. This analysis identified seven modules of co-expressed genes (**Fig. S9A**) capturing a range of expression patterns, with some broadly distributed across cell types and others restricted to specific subsets (**Fig. S9B**).

This variation in cell type specificity is evident in the UMAP coloring (**Fig. 5A**), where some modules are evenly expressed across all cell types, while others are restricted to distinct populations. We quantified the degree of cell type specificity using a cell type specificity index (CTSI), calculated as the standard deviation of the mean module eigengene scores across annotated cell types (**Fig. S9C**). Higher CTSI values indicate stronger restriction to particular lineages. For example, modules 1 and 2 were broadly expressed across nuclei (low CTSI), whereas modules like 7 (enriched in mesoderm and germ cell primordia) and 6 (specific to amnioserosa) showed high CTSI values (**Fig. S9B**).

**Fig. 5.**
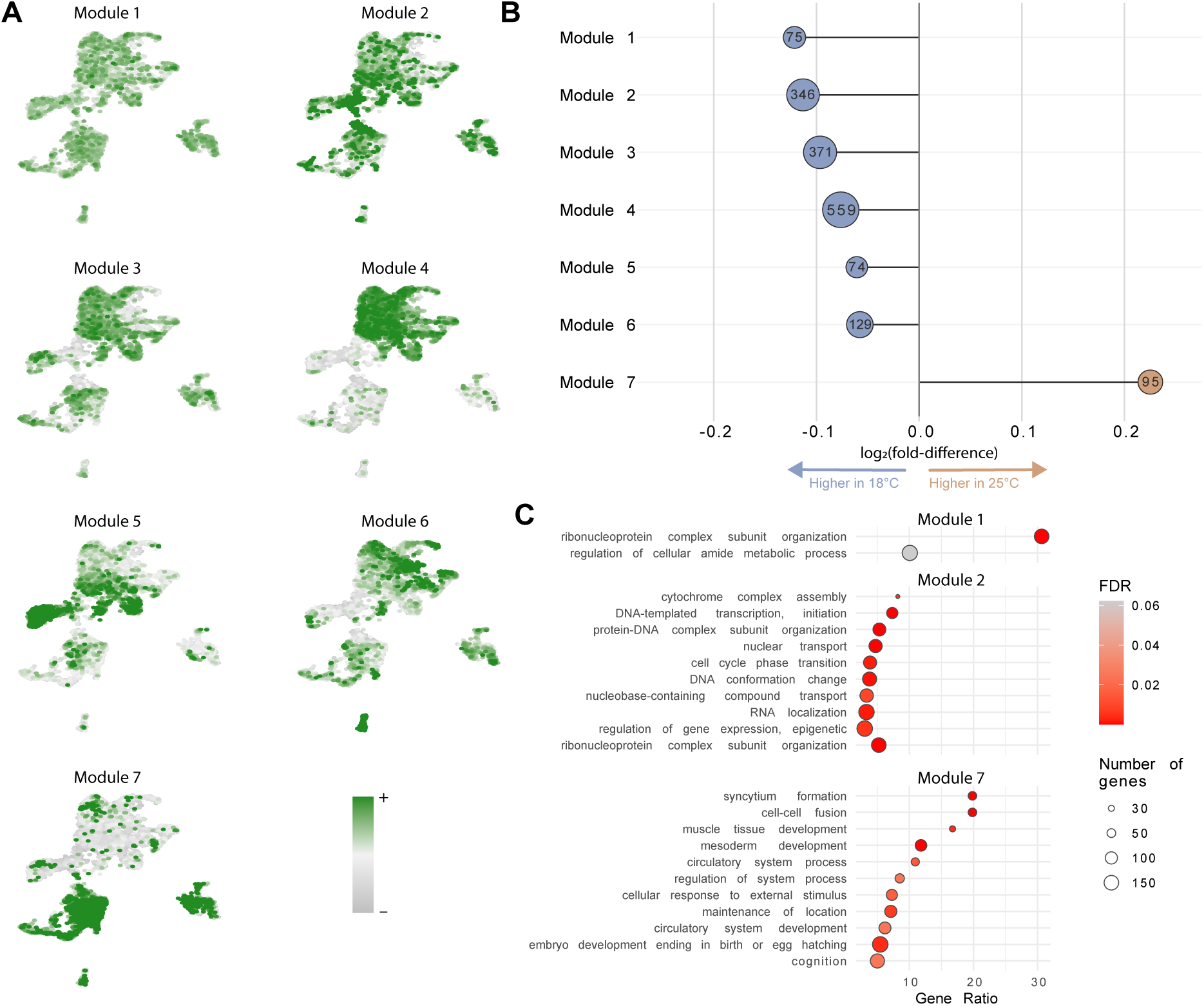
**High-dimensional weighted gene co-expression network analysis (hdWGCNA) on single-nucleus RNA-seq data identifies a few modules of genes that differ in expression following acclimation to different temperatures**. (**A**) UMAPs with each nucleus colored in green by increasing module eigengene value. (**B**) Module eigengene average log2 fold-difference in gene expression between acclimation states for each module. Inset number is the total number of genes per module, with the points sized proportional to module size. Asterisk indicates significance of DME test. (**C**) The top overrepresented GO terms in modules 1, 2, and 7.

Several modules exhibited differential expression between acclimation temperatures, based on changes in average eigengene expression (**Fig. 5B**). Notably, module 7 was significantly upregulated in 25°C-acclimated embryos compared to those acclimated at 18°C (DME test; *p < 0.05*), suggesting a role in temperature-sensitive developmental regulation. Genes in module 7 were enriched for terms related to mesoderm development, muscle tissue development, and embryonic organ development (**Fig. 5C, Fig. S10G**), consistent with expression in specific developmental lineages.

All other modules showed trends of higher expression in cool-acclimated embryos, particularly modules 1 and 2. Functional annotation revealed distinct biological roles for each. Module 1, composed predominantly of ribosomal proteins genes (69 of 75 genes), was enriched for GO terms related to ribonucleoprotein complex organization and amide metabolic processes (**Fig. 5C**). Module 2, containing 346 genes, was enriched for processes including transcription by RNA polymerase III and oxidative phosphorylation (**Fig. 5C, Fig. S10B**). Notably, both modules were broadly expressed across cell types and showed elevated expression in cool-acclimated embryos relative to those at 25°C.

In contrast, modules 3 through 7 displayed greater cell type specificity. Module 3 (371 genes) was expressed primarily in the ectoderm, peripheral nervous system, and ventral nerve cord primordia (**Fig. S9B**). It was enriched for diverse signaling and developmental pathways, including Toll, Smoothened, and Hippo signaling, the MAPK cascade, axis specification, and organ formation (**Fig. S10C**). Module 4, the largest module (559 genes), showed the most polarized expression across cell types, with high expression in ectoderm, peripheral nervous system, ventral nerve cord, foregut & hindgut, and tracheal primordia, and low expression in other cell types (**Fig. S9B**). These genes were highly expressed in 5 cell types. Like module 3, it was enriched for diverse development and signaling pathways (**Fig. S10E**). Module 5 (74 genes) was primarily expressed in the endoderm, ectoderm, and peripheral nervous system primordia. Enriched GO terms included neuroblast differentiation, peripheral nervous system development, and regulation of cell projection organization, consistent with its expression profile (**Fig. S10F**). Module 6 (129 genes) was specific to the amnioserosa (**Fig. S9B**). It was enriched for genes involved in protein processing and cellular organization, including responses to unfolded proteins and ER stress, Golgi vesicle transport, and protein targeting (**Fig. S10D**).

Cell-type-specific hdWGCNA identified 66 gene co-expression modules across all annotated cell types (**Fig. S11**), with 46 modules (70%) trending toward higher expression in 18°C-acclimated embryos. No modules were detected in the amnioserosa, likely due to the small number of nuclei and limited transcriptional heterogeneity, which prevented the network from meeting scale-free criteria. Across cell types, 16 modules were differentially expressed (DMEs; adj. p-value < 0.05) and 13 of these (81%) were upregulated in cool-acclimated embryos. The mesoderm primordium had the highest number of DMEs, with 5 out of 6 modules showing significant acclimation-associated expression differences. We performed GO enrichment analysis to infer the biological functions of the DMEs (**Fig. S12**). While 6 out of 16 DMEs had no significantly enriched terms (FDR < 0.1), the remaining 10 DMEs were enriched for GO terms broadly related to cell-type-specific developmental processes. Notably, this cell-type-specific approach did not recapitulate the broad transcriptional changes observed in modules 1 and 2 of the embryo-wide hdWGCNA, likely because such global patterns are diluted when focusing on individual cell types.

### Cell-type-specific changes in chromatin accessibility and gene expression across acclimation treatments

We next examined cell-type-specific differences in chromatin accessibility and gene expression between acclimation treatments. Before applying a replicate consistency filter, we identified 161 cell-type-specific DARs and 661 DEGs across cell types. We then retained only those DARs and DEGs that showed consistent directionality in both 18°C replicates compared to 25°Ccontrols (**Fig. S13**). All 161 DARs passed this filter, while 50 DEGs were excluded, resulting in 611 high-confidence cell-type-specific DEGs (92.4% of the original set). Concordance between replicates was high. Cell-type-specific DARs showed strong correlation between 18°C replicates (median R^2^ = 0.86, range 0.68-0.91; **Fig. S13A**), and DEGs likewise exhibited high replicate agreement (median R^2^ = 0.89, range 0.61-0.96; **Fig. S13B**), supporting a robust, shared transcriptional and chromatin accessibility response to temperature.

Among the 161 DARs, 100 (62.1%) were more accessible in the 18°C-acclimated embryos, while 61 (37.9%) were more accessible at 25°C (**Fig. 6A**). Similarly, 354 of the 611 (57.9%) were upregulated in18°C-acclimated embryos (**Fig. 6B**). Violin plots illustrate representative examples of these cell-type-specific DEGs (**Fig. S14**). The mesoderm primordium showed the strongest transcriptional response, with 261 DEGs (18.7% of genes tested) and 58 DARs, followed by the peripheral nervous system primordium with 176 DEGs (9.1%) and 19 DARs, all of which were more accessible in the cool-acclimated embryos (**Table S1 and S2**). In contrast, the ventral nerve cord primordium showed greater accessibility in 25°C embryos.

**Fig. 6.**
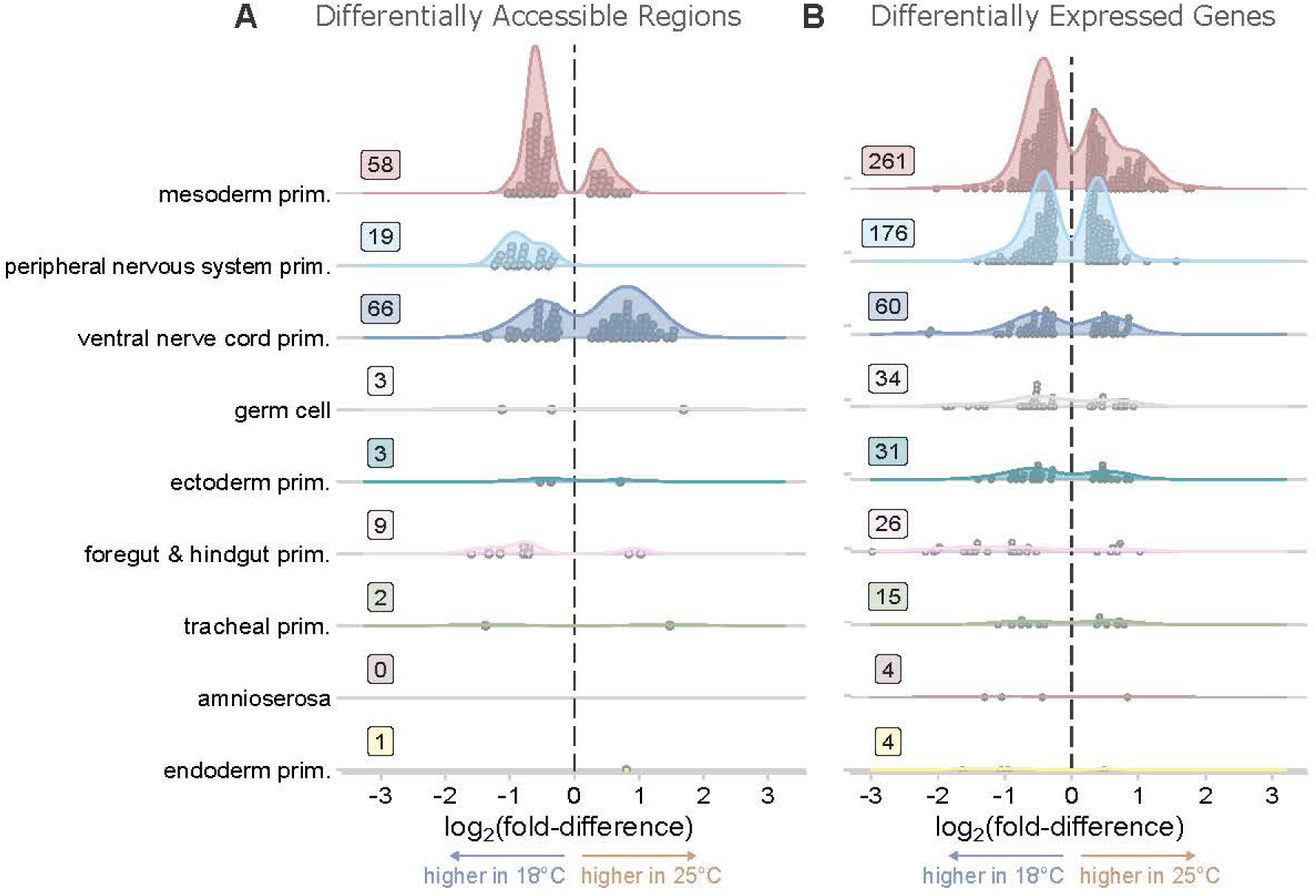
Gene expression and chromatin accessibility differences following embryonic thermal acclimation. Density plots of cell-type-specific log2 fold-differences in accessibility (**A**) and gene expression (**B**). Only significantly differentially accessible regions (DARs) or differentially expressed genes (DEGs) are plotted (adjusted p-value < 0.05). Primordial cell types organized from top to bottom by total number of DEGs. Relative height of the density plots across cell types is proportional to the number of DEGs or DARs. Points represent individual regions or genes. Inset number indicates the total number of DARs or DEGs for each cell-type. Color matches cell-type-specific UMAP in Fig. 2C. Positive log2(fold-difference) values indicate expression or accessibility is greater in the 25°C-acclimated embryos, and negative values indicate greater expression or accessibility in the 18°C cool-acclimated embryos.

At the level of gene expression, the peripheral nervous system showed a mixed response despite all DARs favoring 18°C. The DEGs were nearly evenly split, with 54% (95 genes) upregulated in cool-acclimated embryos. The foregut & hindgut primordium showed increased DARs and DEGs in 18°C embryos, consistent with the *Blimp-1* regulatory network active in these tissues (**Fig. 4B**). Importantly, two cell types with the most DEGs, the mesoderm and peripheral nervous system primordia, also exhibited the most diffGRNs (four and three, respectively), suggesting that cell-type-specific GRNs identified via SCENIC+ likely mediate many of the observed differences in chromatin accessibility and gene expression.

## Discussion

Development in *D. melanogaster* is robust across a broad range of temperatures (*8*, *33*). We asked whether the observed developmental stability reflects passive resilience or arises from active molecular compensation. Overall, our results are consistent with a homeostatic embryonic thermal compensation response to cool acclimation, whereby organisms actively increase metabolic activity to overcome slower kinetics at cooler temperatures. In cool-acclimated embryos, WGCNA revealed global upregulation of gene modules enriched for ribosomal proteins, transcription factors, and oxidative phosphorylation genes. Gene regulatory network analysis showed that six of the eight activator diffGRNs exhibited increased transcription factor expression, motif accessibility, and target gene expression in the cool-acclimated condition. Additionally, peaks containing the *Zelda* pioneer factor motif were globally more accessible, suggesting enhanced capacity for zygotic genome activation (*62–64*).

Thermal compensation has been observed across a broad range of taxa; it is worth noting, however, that previous work has not described compensation in the context of developmental robustness and typically involved much longer acclimation periods (*65– 71*). Evidence of complete thermal compensation — *i.e.*, where cold-acclimated organisms exhibit the same biological rates in the cold as organisms acclimated to warmer environments — has not been documented (*72*). Thus, as cool-acclimated *D. melanogaster* embryos develop more slowly, they likely achieve only partial thermal compensation (*8*). While we did not directly measure metabolic or transcriptional rates, the observed shifts in chromatin accessibility and gene expression in cool-acclimated embryos suggest partial compensation of biological rates.

However, these compensatory molecular responses may incur trade-offs in acute heat tolerance. Stage 11 embryos exhibited rapid thermal plasticity in heat tolerance, adjusting within hours to match their environmental temperature. This effect was most pronounced at low rearing temperatures, with a nearly 50% decline in survival following acute heat stress for 18°C embryos relative to 25°C. Elevated protein synthesis can increase the risk of protein misfolding and denaturation (*73*, *74*), while increased oxidative phosphorylation may elevate reactive oxygen species production and oxidative damage (*75*, *76*). Together, these coordinated transcriptional and chromatin-level changes likely buffer embryonic development in the cold but may impose metabolic costs that reduce resilience to heat stress — an underappreciated consequence of maintaining developmental robustness.

Regulatory changes concentrated in specific cell types may also contribute to shifts in thermal performance. GRN shifts were restricted to the mesoderm, gut, and nervous system primordia, cell types that also exhibited the highest number of DARs and DEGs, underscoring the importance of localized regulatory mechanisms. In adult *Drosophila*, these same tissues play important roles in regulating the thermal stress response. Ion homeostasis in gut and muscle is critical for cold tolerance (*77–80*), the heart contributes to heat tolerance (*81*), the larval gut mitigates heat-induced cellular damage (*82*), and neural function is particularly sensitive to elevated temperatures (*83*). While embryonic primordia differ from their differentiated counterparts, they share functional regulators such as *Blimp-1, Mef2,* and *ab,* which are involved in gut, muscle, and neural development and maintain roles in ion transport, neuromuscular function, and contractility in mature tissues (*84–86*). These findings suggest that gut, muscle, and nervous system primordia may play early, active roles in shaping thermal responses during embryogenesis, an avenue for future investigation.

Follow-up research is required to quantify the extent of thermal compensation in *D. melanogaster* embryos (*e.g.*, using respirometry to measure metabolic rates or nuclear run-on assay to measure transcriptional activity). Additionally, future studies could directly test the role of *Zelda* and other pioneer factors in chromatin remodeling under cool acclimation by measuring protein–DNA interactions (ChIP-seq) across temperatures. Our current data cannot determine whether the increased accessibility of ZLD-target-motifs reflects increased ZLD activity, as *Zelda* is maternally deposited and translated in the hour after fertilization (*26*, *87*). However, because ZLD is essential for zygotic genome activation, it is notable that the majority of ZLD-target-motifs were more accessible in the 18°C-acclimated embryos. Although all embryos were hand-staged to the same developmental time point, they were undergoing rapid cell specification (*48*). Thus, finer temporal resolution and deeper sequencing could reveal additional regulatory changes.

Thermal acclimation in *D. melanogaster* embryos occurs remarkably fast. Whereas later life-stages require days or weeks to acclimate (*65–71*), *Drosophila* embryos responded within hours, highlighting potential life-stage-specific differences in plasticity. Moreover, beneficial thermal acclimation (*i.e.*, a positive relationship between acclimation temperature and upper thermal limits) is not universal. Some taxa show no response or even reduced heat tolerance at higher temperatures (*88–90*), which may reflect increased reliance on behavioral thermoregulation, relaxed selection for physiological plasticity (*91*), or species and life stages living near their upper thermal limits, limiting further plasticity (*23*, *88*). Given their immobility, narrow thermal window, and the rapid timescale of potential temperature variability (*18*), even small shifts in embryonic heat limits could be critical for survival in natural systems (*17*).

These findings suggest that the classic view of environmental robustness in development should be refined to account for underlying molecular plasticity. While morphological outcomes are often canalized (*14*, *92*, *93*), this may come at a cost for other traits such as heat tolerance (*94*, *95*), which appeared reduced by homeostatic plasticity. Both morphological and physiological traits are essential for survival (*96–98*), but selection for morphological canalization may favor homeostatic mechanisms that reduce physiological flexibility. We propose that compensatory responses to cold, such as increased metabolic activity, may induce cellular stress under subsequent heat exposure, leading to a trade-off. This work highlights the complex interplay between developmental robustness and physiological plasticity in shaping species’ resilience to climate variability.

## Materials and Methods

### Fly care

We raised Canton-S flies (Canton, OH, USA) on standard yeast, cornmeal, and molasses food. We maintained stocks at a standard density of approximately 50 to 100 flies per vial (95mm × 25mm, Genesee Scientific) in an incubator (DR-36VL, Percival Scientific Inc.) at 25°C and 55% relative humidity on a 12 hour:12 hour light:dark cycle. Flies were reared under these conditions for several generations prior to experimentation.

### Acclimation and assessing heat tolerance

At least 48 hours prior to embryo collections, we transferred approximately 100 mating pairs of ∼1-2 day-old adult Canton-S flies into small fly cages (Genesee Scientific) with grape juice agar plates (60 × 15 mm) with yeast paste. Fly cages remained in the 25°C incubator. Immediately prior to each experimental collection, we conducted two successive one-hour pre-lays to give the flies fresh egg-laying substrate and reduce the incidence of egg retention. With a fresh agar and yeast plate, we collected embryos for one hour and then transferred the embryos to an incubator at the acclimation treatment temperature (18°C, 25°C, or 30°C). Embryos developed for a certain amount of time based on the expected development rate at their acclimation temperature. We estimated the development times based on a *Q*_10_ of 2.2 (*8*). We subsequently validated these estimated development times by visualizing Bownes’ stage 11 embryos in each acclimation condition under a light microscope (M80, Leica). Stage 11 embryos were defined as embryos displaying full germ band extension towards the anterior. Control embryos developed at 25°C for 5 hours after egg collection, cool-acclimated embryos developed for 9 hours at 18°C, and the warm-acclimated embryos developed at 30°C for 3 hours.

To assess acute heat tolerance following acclimation, we wrapped egg plates in Parafilm and submerged them in a 38.75°C water bath (A24B, Thermo Scientific) for 45 minutes. Following the heat shock, we placed individual embryos into four-by-five grids of twenty total eggs (Total number of eggs per acclimation temperature: 520, 1272, 1145 for 18°C, 25°C, and 30°C respectively). The egg plates with gridded embryos recovered from heat shock at 25°C, and we scored hatching success approximately 48 hours after egg laying by visualizing hatching under a dissecting microscope. To contrast heat tolerance among acclimation treatments, we conducted a quasi-binomial logistic regression generalized linear model in R on the proportion of eggs hatched after acute heat stress. Later, to test the pairwise differences between groups, we conducted a Tukey-adjusted pairwise Wald test on estimated marginal means.

### Embryo acclimation and nuclei extraction for sequencing

Following the heat tolerance results, we chose to move forward with the control 18°C-and 25°C-acclimated embryos for Chromium Single Cell Multiome ATAC + Gene Expression (10x Genomics). We collected and acclimated embryos as above, but immediately following acclimation, we washed and dechorionated the embryos. For each sample, approximately 50 stage 11 embryos were hand-selected and collected with a paintbrush into a 1.5 mL DNA LoBind tube (Eppendorf). We added 100 µL of cryobuffer (90% FBS and 10% DMSO), placed the samples in an isopropanol cryochamber (Cryo-Safe™ Cooler-1°C Freeze Control, Bel-Art Products) and then froze them at –80°C for less than three weeks until nuclei extraction.

Nuclei extraction was completed as described in Albright *et al.* (2022) (*112*). In detail, we thawed the cryofrozen embryo samples and subsequently washed the embryos with 250 µL 1× PBS. We centrifuged (Sorvall ST89, Thermo Scientific) the washed samples at 500 RCF for three minutes at 4°C. To the resulting washed embryo pellet, we added 600 µL of lysis buffer (10 mM Tris-HCl, pH 7.5, 10 mM NaCl, 3 mM MgCl∼2∼, 0.1% IGEPAL, 1% BSA, 1 mM DTT and 1 U/µL RNase Inhibitor – RNaseOUT™ Recombinant Ribonuclease Inhibitor, Thermo Fisher – prepared with nuclease free H_2_O, and then we transferred the sample to a 1 mL Dounce homogenizer (KIMBLE® KONTES®, Sigma). We homogenized the embryos on ice with the loose pestle for ten passes and then with the tight pestle for an additional ten passes. To decrease loss of sample, we rinsed the pestles with 100 µL of lysis buffer following removal from the sample. We filtered each 800 µL homogenate with a 40 µm Mini-Strainer (pluriSelect) into separate fresh 1.5 mL LoBind tubes. We centrifuged the samples at 900 RCF for five minutes at 4°C. After discarding the supernatant, we washed the resulting nuclei pellet with 500 µL wash buffer (identical to the Lysis buffer but missing IGEPAL) and centrifuged again at 900 RCF for five minutes at 4°C. Finally, we resuspended the washed nuclei pellet in 20 µL 1× nuclei buffer (10x Genomics) with 1 mM DTT and 1 U/µL RNase Inhibitor.

To assess the quality and concentration of each sample, we visualized an aliquot of isolated nuclei at one-tenth dilution in 1× PBS and 0.1 mg/mL DAPI on a confocal microscope (Nikon ECLIPSE Ti2). We confirmed nuclei quality by looking for the absence of nuclear blebbing at 600× magnification. All visualized nuclei were high-quality or mostly intact with minor evidence of blebbing as defined by 10x Genomics. We estimated nuclei concentration by loading 10 µL of one-tenth diluted sample onto a hemocytometer (Bright-Line™, American Optical) and visualized nuclei with DAPI fluorescence on the confocal microscope at 100× magnification.

Prior to proceeding to 10x Genomics Chromium Single Cell Multiome ATAC + Gene Expression sequencing library preparation, we diluted each sample to an estimated concentration of 2,000 nuclei per µL with 1× nuclei buffer.

### Library preparation and sequencing

Vermont Integrative Genomics Resource (VIGR) conducted the ATAC and GEX library preparation following the 10x Genomics protocol (Chromium Next GEM Multiome ATAC + Gene Expression User Guide, Document Number CG000338, Rev F, August 2022). We targeted 4,000 total nuclei per sample. Briefly, each sample of nuclei was transposed and then loaded onto a Chromium Next GEM Chip J to generate GEMs and barcode individual nuclei. After the GEMs are broken down, barcoded transposed DNA and cDNA are amplified with PCR. Finally, we constructed the individual ATAC and Gene Expression libraries and sequenced them on a rapid-run flow cell on an Illumina HiSeq 2500.

### Sequencing analysis

We conducted all sequence processing on the Vermont Advanced Computing Core (VACC). Sequencing reads were demultiplexed and analyzed with the Cell Ranger ARC, version 2.0.2 (10x Genomics) pipeline as recommended. In detail, a *D. melanogaster* reference package was created following 10x Genomics recommendations. The FASTA (BDGP6.32; Drosophila_melanogaster.BDGP6.32.dna.toplevel.fa.gz) and GTF (BDGP6.32.109; Drosophila_melanogaster.BDGP6.32.109.gtf.gz) files were downloaded from Ensembl database. Prior to creating the reference package, the GTF file was filtered so that the reference was restricted to protein coding, long non-coding RNA, antisense, and immune-related genes, the same filter criteria used by 10x Genomics to create the human and mouse references.

Then, the Cell Ranger ARC (version 2.0.2) pipeline was run on all samples to trim, align, and map reads, and then create count matrices for gene expression and chromatin accessibility peaks. Samples were then aggregated together into a single file for each data type and then the files were imported into R version 4.2.2 (*99*) and analyzed using Seurat version 4.4.0 (*100*) and Signac 1.12.0 (*101*). Subsequently, we used python version 3.8.18 to model *cis*-regulatory topics using pycisTopic version 1.0.3 (*102*), then transcription factor motif enrichment using pycisTarget version 1.0.3, and finally identified enhancer-driven gene regulatory networks using SCENIC+ version 1.0.1 (*61*). All code used for the analysis is available on GitHub at https://github.com/tsoleary/heater.

### Quality control filtering

Single GEM barcodes were filtered to high quality nuclei based on several criteria: (i) a low-count threshold for both the ATAC and RNA libraries (≥ 800 and ≥ 200 counts per barcode respectively), (ii) a maximum cut-off for percentage of reads mapped to mitochondrial transcripts (< 5%) and ribosomal genes (< 25%), (iii) putative multiplets were identified using AMULET for the ATAC-seq libraries (*103*) and DoubletFinder for the RNA-seq libraries (*104*). Finally, all barcodes in clusters with > 15% putative multiplets were removed. This filtering left 10,390 high-quality nuclei for all downstream analyses.

Due to the high cell-cycle heterogeneity expected in developmental tissue, genes known to change across cell cycle phases were regressed out of the snRNA data (*100*). We called chromatin accessibility peaks using MACS2 version 2.2.7.1 (*105*), and analyzed the RNA and ATAC data in Seurat according to current recommendations (*101*, *106*).

### Clustering and annotating cell types

We identified clusters from the joint ATAC and RNA data using the FindMultiModalNeighbors and FindClusters function in Seurat. We tested multiple clustering resolutions ranging from 0.1 to 1, and settled on a resolution of 0.7 (Fig. S3). We then used the gene expression information to identify marker genes for each cluster using the FindConservedMarkers functions. The clusters were assigned annotated cell types by conducting a Fisher’s exact test for enrichment on marker genes annotated by the Berkeley *Drosophila* Genome Project RNA in situ hybridization database (*28*).

### Differentially enriched motifs in pseudobulk DARs

We tested for differentially enriched motifs using the FindMotifs function in Seurat. *D. melanogaster* motifs were identified using the transcription factor binding profiles from JASPAR 2022 (*107*).

### Cell-type-specific differential expression and accessibility testing

Using Seurat, we ran differential accessibility and expression testing between acclimation treatments but within cell types using MAST (*49*) within the FindMarkers function in Seurat. In addition to a Bonferroni adjusted p-value less than 0.05, differentially accessible regions (DARs) or differentially expressed genes (DEGs) needed to be present in greater than 10% of nuclei tested and a have minimum absolute of log_2_ fold-difference of 0.25. To ensure that the signal of differential expression and accessibility was a shared response to acclimation, we added an additional criterion to our DEG & DAR calls by conducting a sample exclusion analysis. In the sample exclusion analysis, we subsetted each individual 18°C-acclimated replicate and tested that sample against the 25°C-acclimated control nuclei and only kept DEGs & DARs that were consistent across samples.

### Weighted gene co-expression network analysis (hdWGCNA)

We conducted a high dimensional weighted gene co-expression network analysis with hdWGCNA (*108*); version 0.2.27) on the single-nucleus RNA data to identify modules of genes that are expressed in similar patterns as well as modules that are different between acclimation conditions. According to package recommendations, we only included genes that were detected in greater than 5% of nuclei. Differential module eigengenes between acclimation conditions were identified using the FindDMEs function using MAST similar to the individual DEG and DAR testing as described above. Additionally, due to the expectation that each individual cell type is likely to have its own networks of co-expressed gene modules, we also subsetted the data for each cell type and conducted the same hdWGCNA analysis on all nine primordial cell types.

### Identifying enhancer-driven gene regulatory networks using SCENIC+

SCENIC+ (version 1.0.1) was used to identify gene regulatory networks (GRNs). Using the single-nucleus ATAC-seq data, cis-regulatory topics were modeled using cisTopic (*102*), and cisTarget was then used for motif enrichment and identifying transcription factor cistromes. SCENIC+ was used to identify cell-type-specific enhancer-driven gene regulatory networks (GRNs) by using the linked gene expression data from the single-nucleus RNA-seq (*61*). We tested for differential GRNs (diffGRNs) that were altered by acclimation using a three-part test: (i) the transcription factor is differentially expressed within the cell-type (DEGs MAST; |lfc| > 0.25, adjusted p-value < 0.05, percentage of nuclei > 10%), (ii) the distribution of log_2_ fold-differences of the regions containing the target motif are shifted away from zero in the expected direction (one-sample t-test; p < 0.05), and (iii) the distribution of log_2_ fold-differences of the corresponding target genes are shifted in the same way (one-sample t-test; p < 0.05). This method tests for GRNs that have changed their transcription factor expression, target motif accessibility, and target gene expression as a result of acclimation.

## Acknowledgments

We thank Kristi Montooth, Colin Meiklejohn, and Princess Rodriguez for helpful discussions.

## Author contributions

Conceptualization: TSO, SF, SHC, BLL

Methodology: TSO, SF, SHC, BLL

Investigation: TSO, EEM, ST

Formal analysis: TSO, JRB, SF, BLL

Supervision: BLL, SHC, SF

Writing — original draft: TSO, BLL

Writing — review & editing: TSO, BLL, SF, SHC

## Data and materials availability

Multiome sequencing data is available on the Sequence Read Archive. All code used for analysis and data visualization available on GitHub (https://github.com/tsoleary/heater).

## Supplementary Materials

**Fig. S1.**
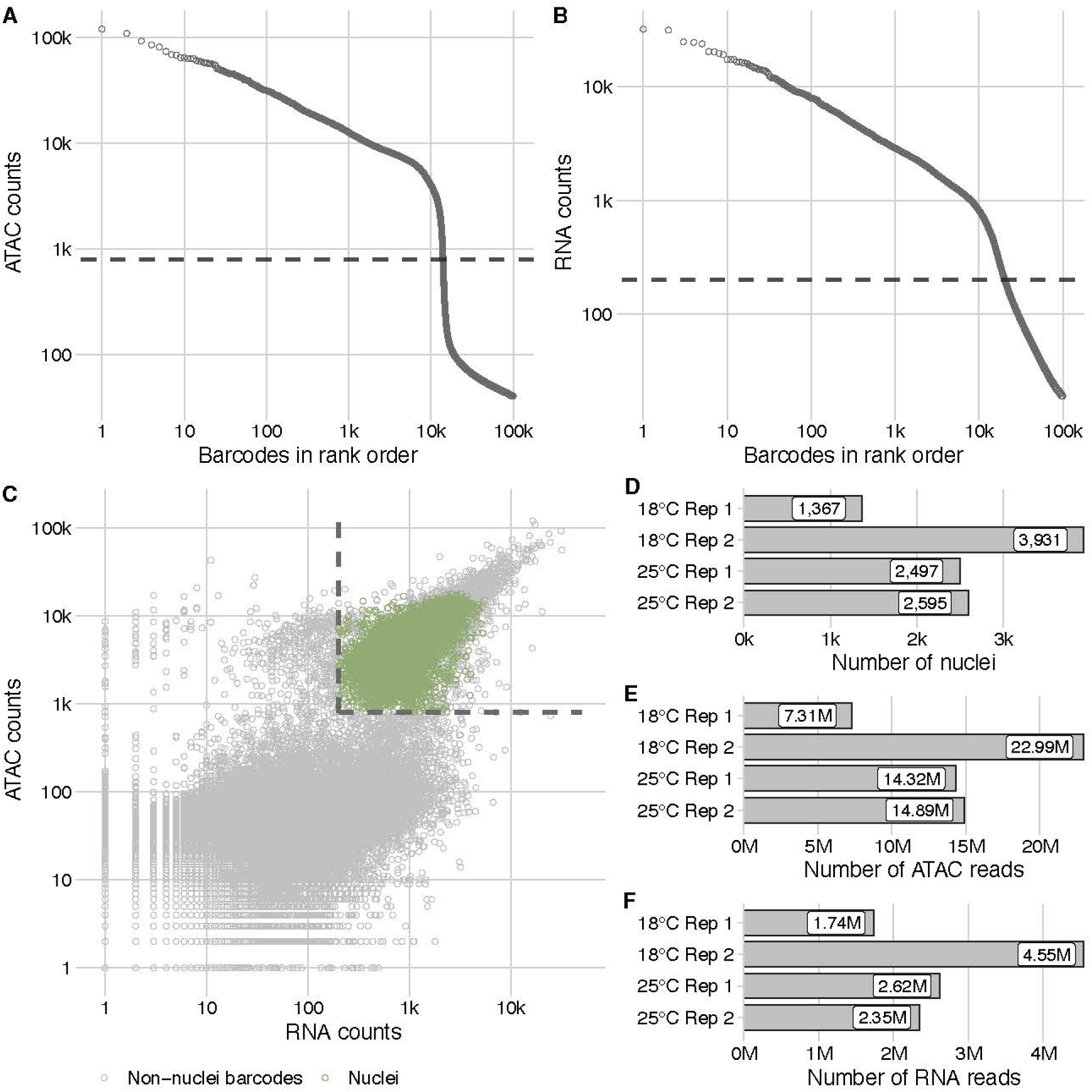
Overall multiome sequencing quality control metrics for cell filtering and total read depth. (**A**) Scatter plot of the number of ATAC reads per barcode in rank order from highest to lowest, transformed onto a log scale for both axes. Showing only the top hundred thousand barcodes. Dashed line represents the minimum cut off for filtering (800 reads). (**B**) Scatter plot of the number of RNA reads per barcode in rank order from highest to lowest, transformed onto a log scale for both axes. Showing only the top hundred thousand barcodes. Dashed line represents the minimum cut off for filtering (200 reads). (**C**) Scatter plot of ATAC and RNA counts for all non-zero barcodes. Green dots represent high quality nuclei after filtering. Dashed lines represent the minimum cutoff for the number of RNA and ATAC reads. (**D**) Bar plot of the number of final high-quality nuclei from each sample. (**E**) Total number of ATAC-seq reads from each sample. (**F**) Total number of RNA-seq reads from each sample.

**Fig. S2.**
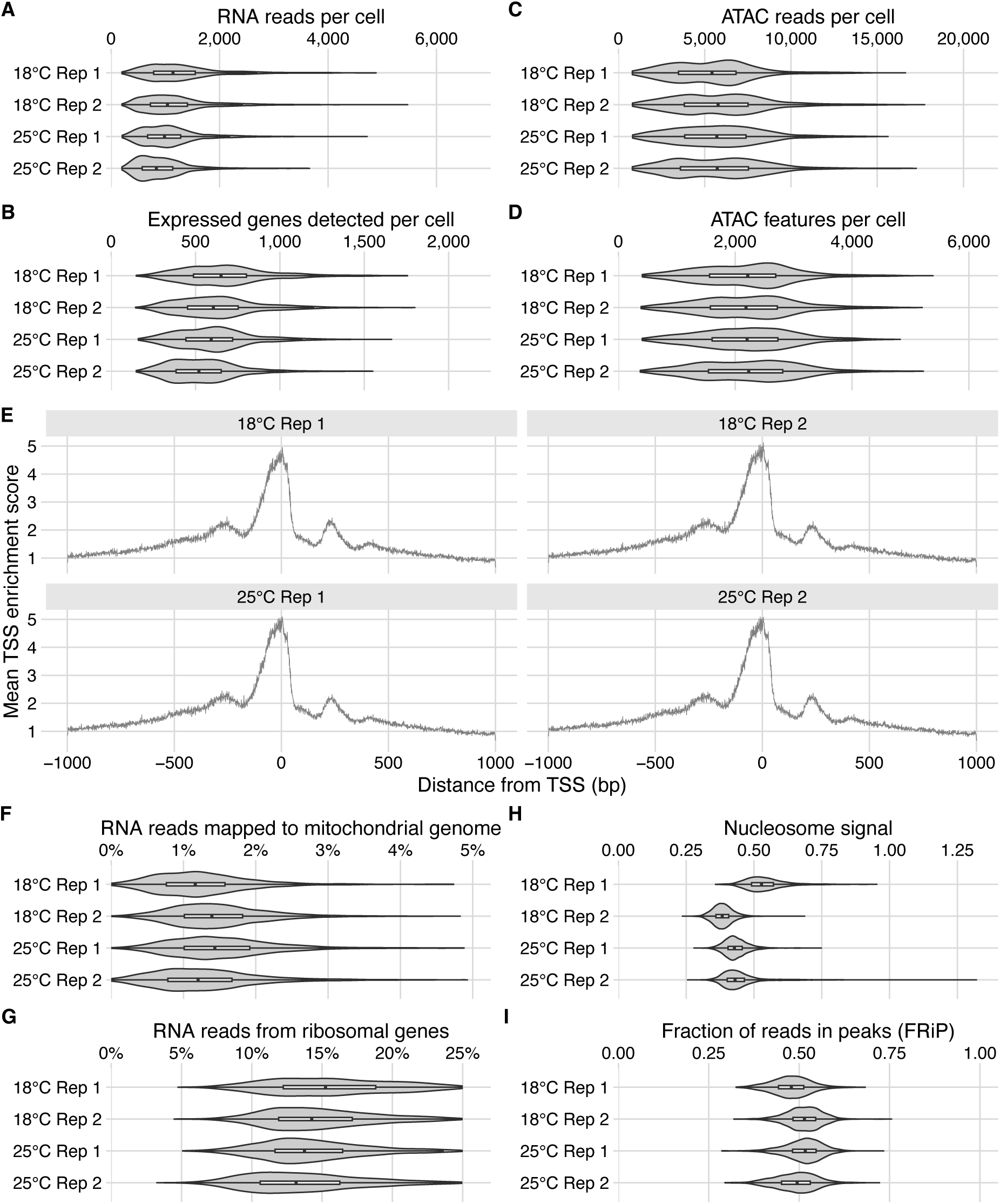
Per nucleus quality control metrics after filtering. (**A**) Total number of RNA reads. (**B**) Number of genes detected. (**C**) Total number of ATAC reads. (**D**) Total number of ATAC features. (**E**) TSS enrichment. (**F**) Percent RNA reads mapped to the mitochondrial genome. (**G**) Percent RNA reads mapped to ribosomal genes. (**H**) Nucleosome signal as defined by the ratio of mononucleosome fragments (between 147 bp and 294 bp) to nucleosome-free fragments (< 147 bp). (**I**) Fraction of ATAC reads in peaks (FRiP).

**Fig. S3.**
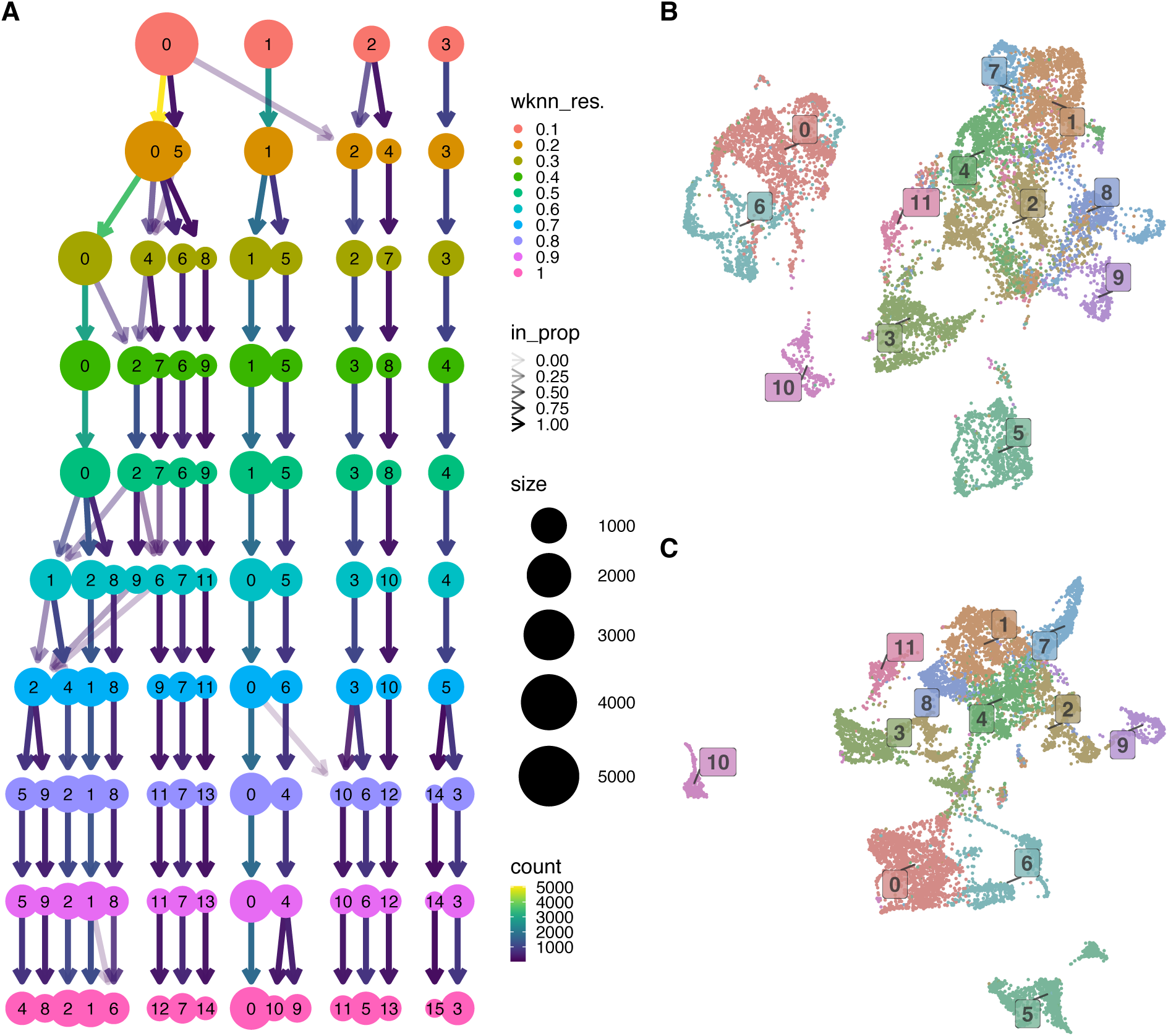
Joint clustering at different resolutions alongside RNA-only and ATAC-only UMAPs with the final clustering resolution visualized. (**A**) Clustree representation of the Seurat clusters at different resolutions ranging from 0.1 to 1.0, with a resolution of 0.7 being the final chosen resolution. UMAP representation of all nuclei using only the ATAC-seq data (**B**), and only the RNA-seq data (**C**) colored by Seurat clusters of the joint ATAC & RNA clustering.

**Fig. S4.**
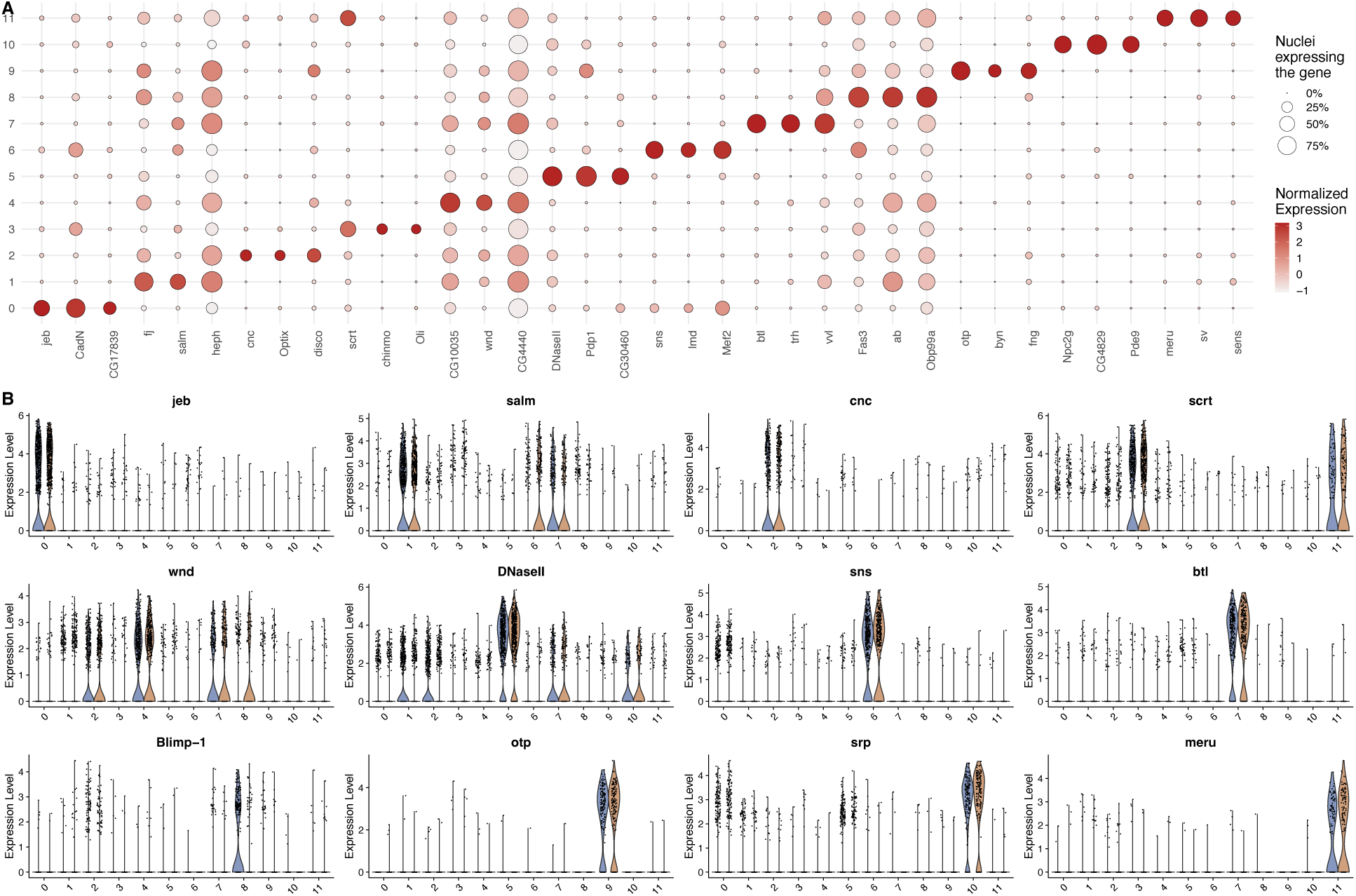
Top marker genes from RNA-seq data for each cluster show cluster-specific expression patterns. (**A**) Dot plot of the top three marker genes from each cluster by lowest p-value. Colored red by normalized expression. Dots are sized by the percent of nuclei within a cluster where that marker gene was detected. (**B**) Violin plots of representative marker genes from each of the 12 clusters. Each cluster is split by acclimation treatment indicated by color: on the left, blue represents 18°C-acclimated embryos, and on the right, orange represents 25°C-acclimated embryos.

**Fig. S5.**
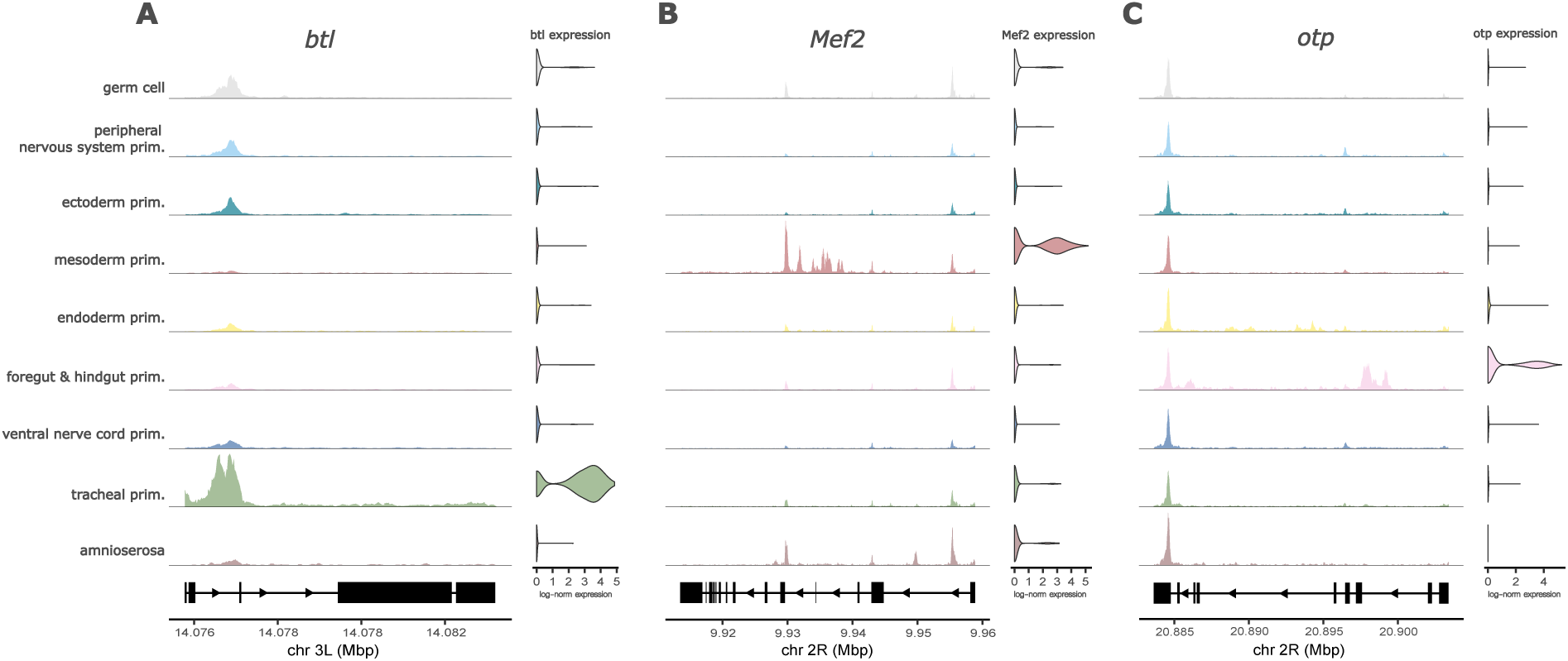
ATAC-seq coverage plots (left panel) of peaks linked to marker genes show that the cell-type-specific accessibility matches the cell-type specific expression (violin plots on the right panel) for three representative marker genes *btl* (**A**), *Mef2* (**B**), and *otp* (**C**), in the tracheal, mesoderm, and foregut & hindgut primordia respectively.

**Fig. S6.**
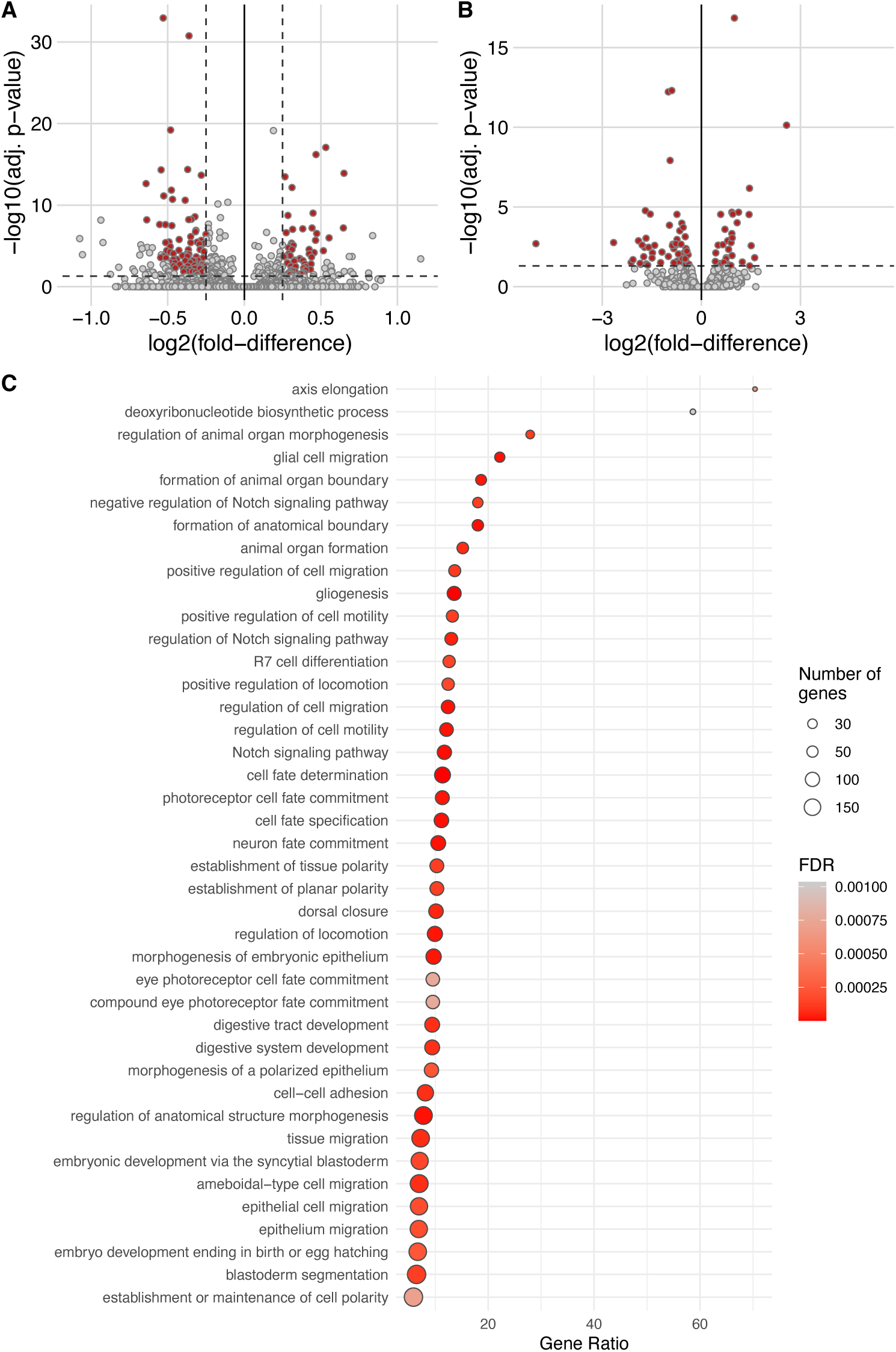
Results from the pseudobulk analysis. (**A**) A volcano plot of the differences in accessibility (ATAC-seq) between acclimation treatments with differentially accessible regions marked in red. (**B**) A volcano plot of the differences in expression (RNA-seq) between acclimation treatments with differentially accessible genes marked in red. (**C**) A dot plot of the 41 over-represented GO categories in the genes linked to the 33 *Zelda*-target-motif containing DARs.

**Fig. S7.**
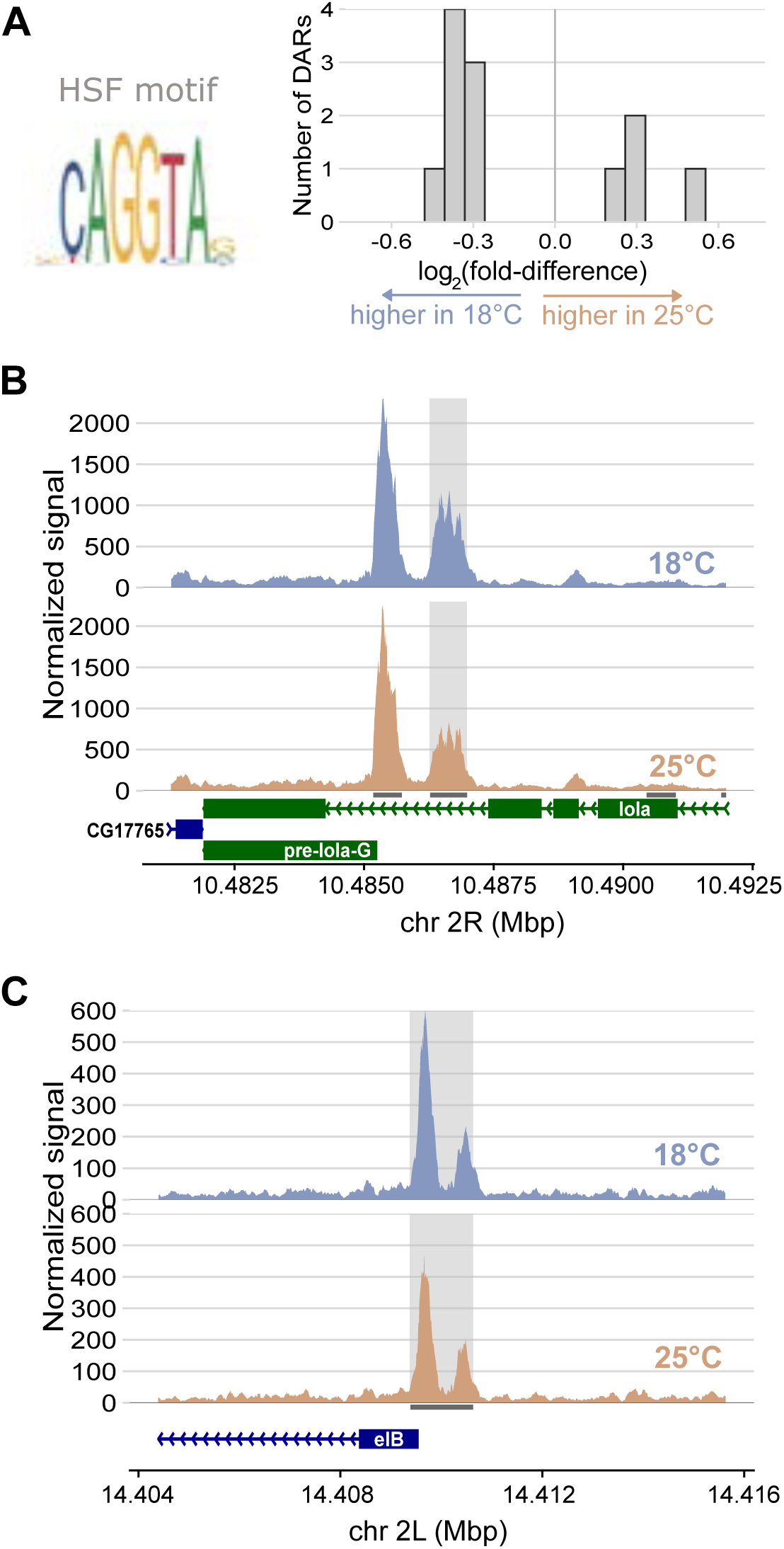
(**A**) Histogram of log2 fold-difference between acclimation treatments of the 12 pseudobulk DARs that contain the HSF-target motif. Pseudobulk ATAC-seq coverage plots for three representative HSF-target motif containing DARs (**B-C**). Within each panel containing the coverage plots, the top trace, colored blue, represents the accessibility of the 18°C-acclimated embryos, and the bottom trace, in orange, represents the 25°C-acclimated embryos. Grey shading indicates the peak regions with differential accessibility between acclimations.

**Fig. S8.**
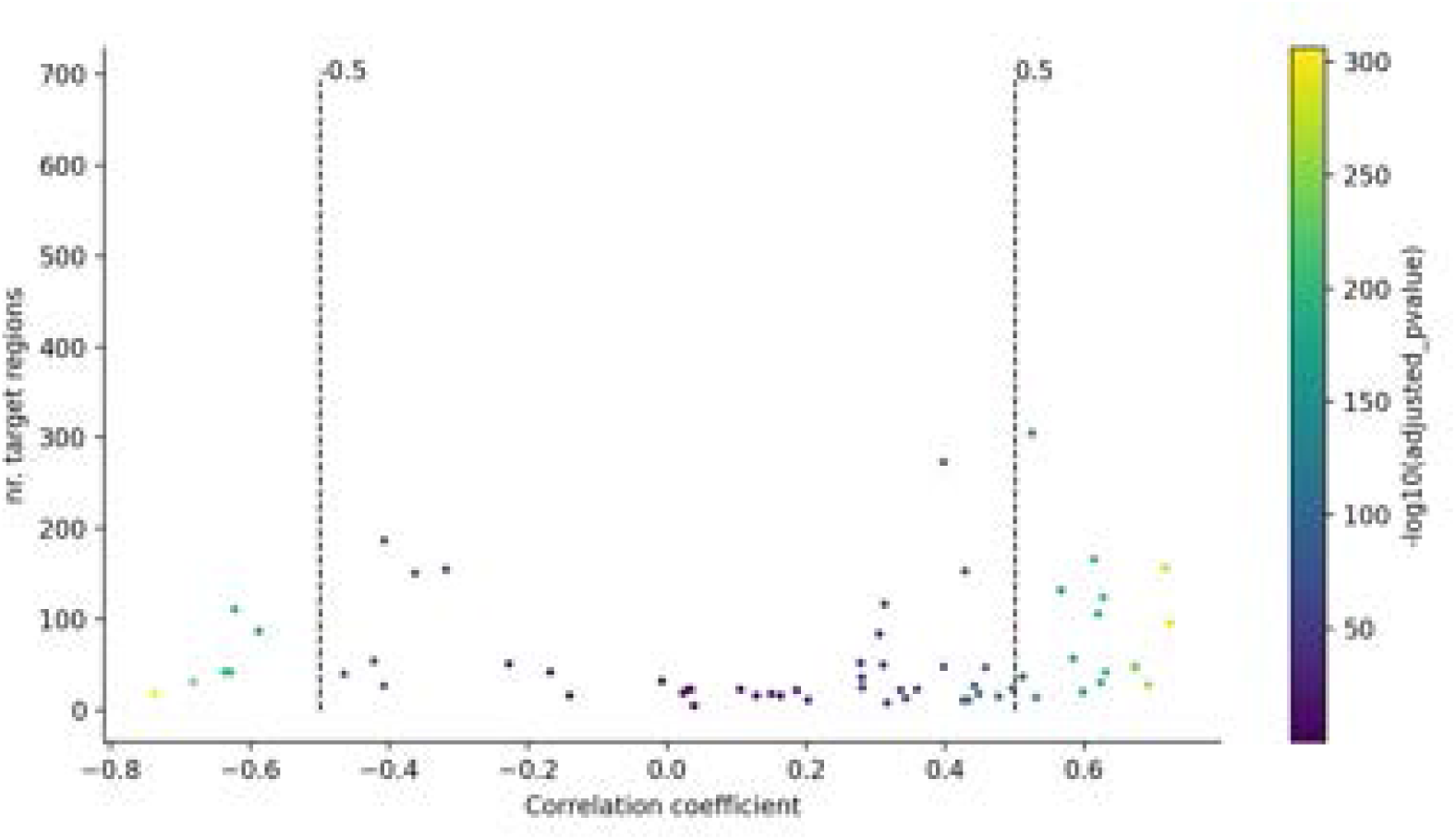
41 GRNs identified in the SCENIC+ pipeline with the highest correlation coefficients (|rho| > 0.5) were selected for downstream analysis. Each point represents an individual GRN. Points are colored by adjusted p-value. The values along the vertical axis represent the number of target regions in each individual GRN and the values of the horizontal axis represent the correlation coefficient (rho) calculated during eGRN inference.

**Fig. S9.**
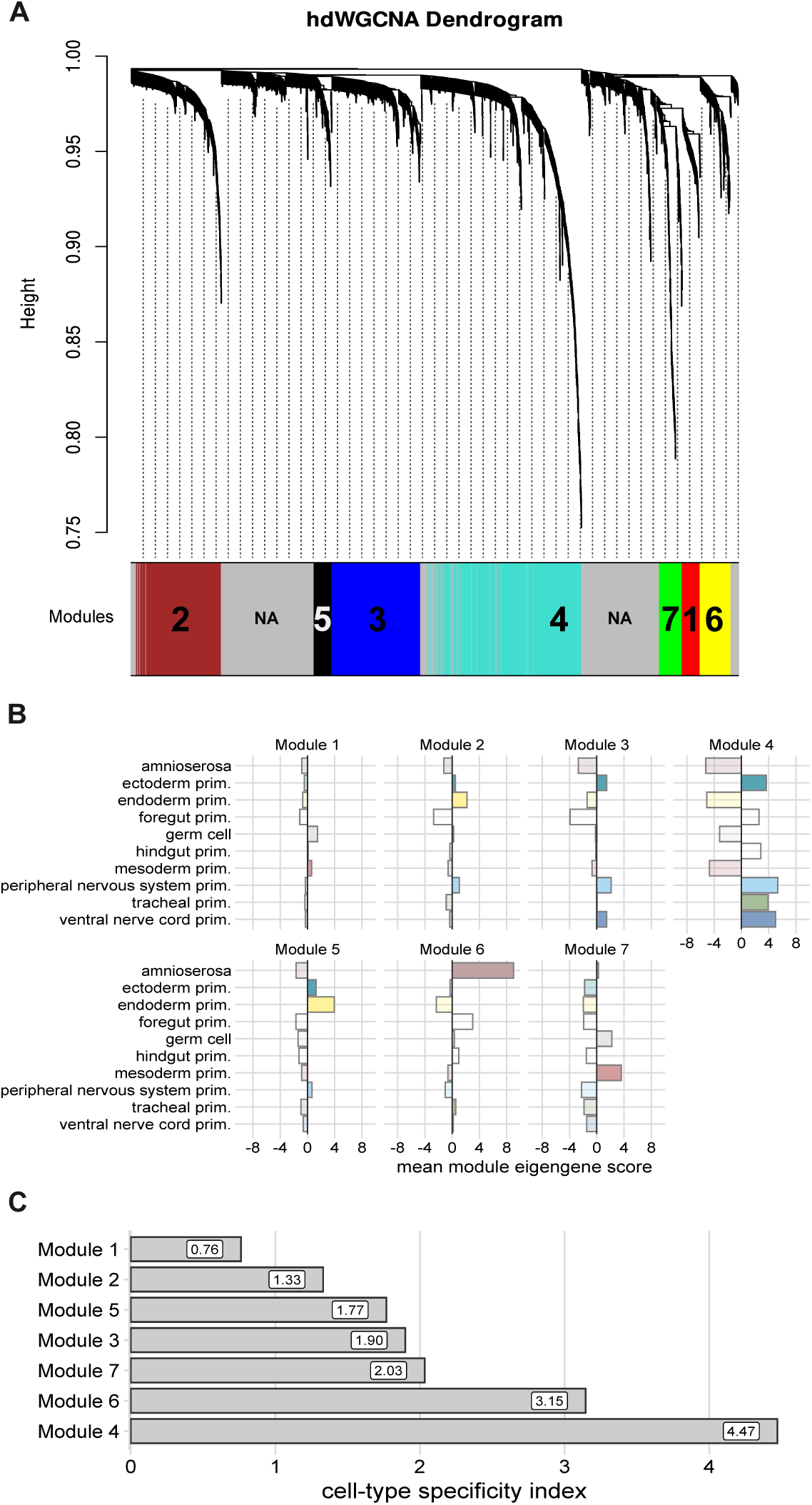
hdWGCNA module dendrogram and the cell-type specificity of modules. (**A**) hdWGCNA identified seven modules of co-expressed genes. (**B**) Mean module eigengene score for each cell-type. (**C**) Cell-type specificity index (CTSI) for each module was calculated as the standard deviation of the mean cell-type-specific module eigengene scores. Low values represent modules that are expressed evenly across tissue types (*i.e.,* the genes within the modules with low CTSI are expressed generally across cell types – low cell-type specificity). High values indicate that the genes within the module are more specific to certain cell types (*i.e.,* genes within the modules with high CTSI are expressed predominantly within only a subset of cell types – high cell-type-specificity).

**Fig. S10.**
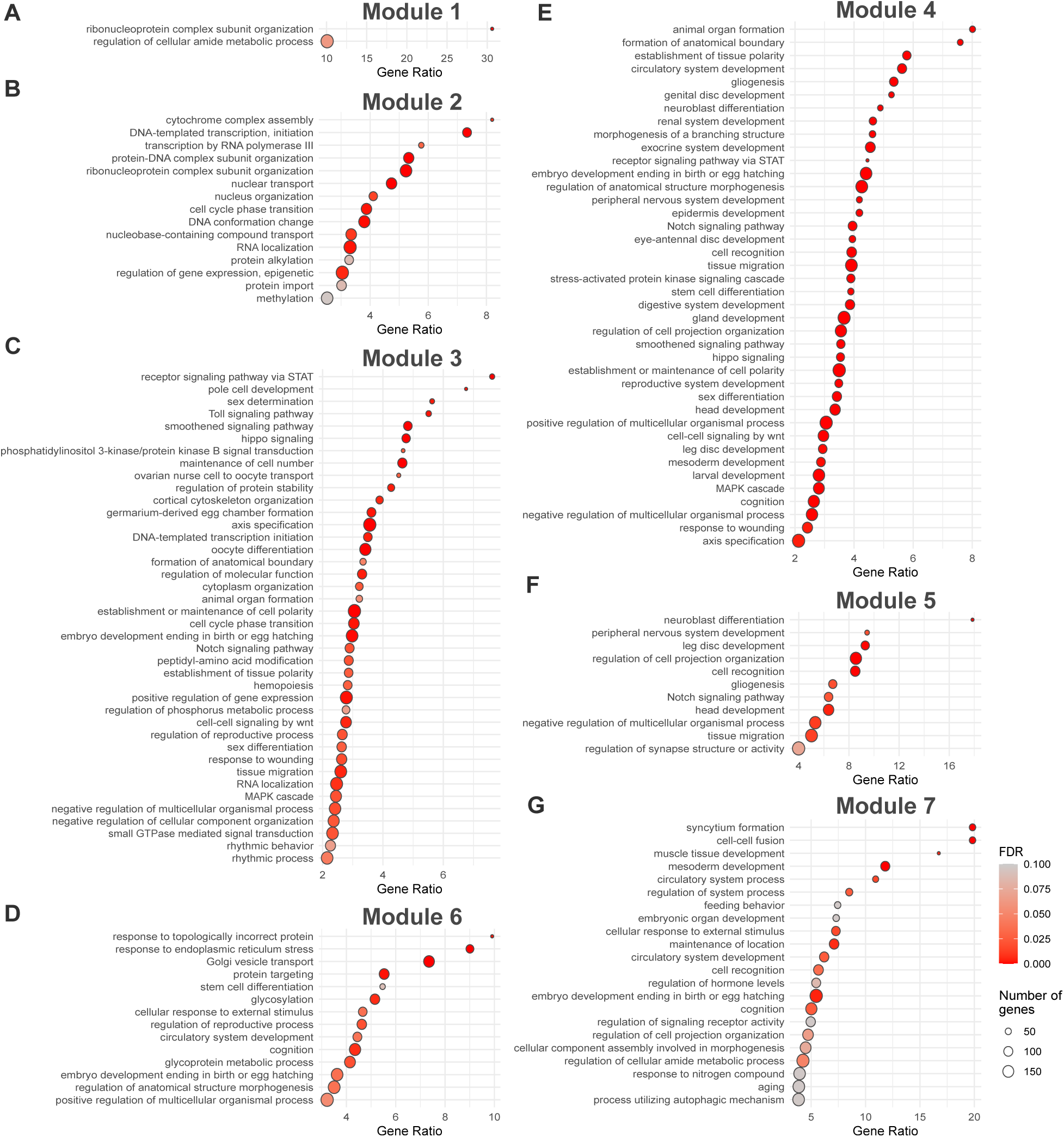
GO terms of overrepresentation analysis of all seven modules of co-expressed genes identified by hdWGCNA. Points are sized by the number of genes in the GO set and colored by the FDR. Horizontal axis indicates the gene ratio (i.e., the number of DEGs present in a given GO set divided by the number of DEGs expected in that GO set by random chance, based on the background distribution of all expressed genes). All significantly overrepresented GO terms are included (FDR < 0.1).

**Fig. S11.**
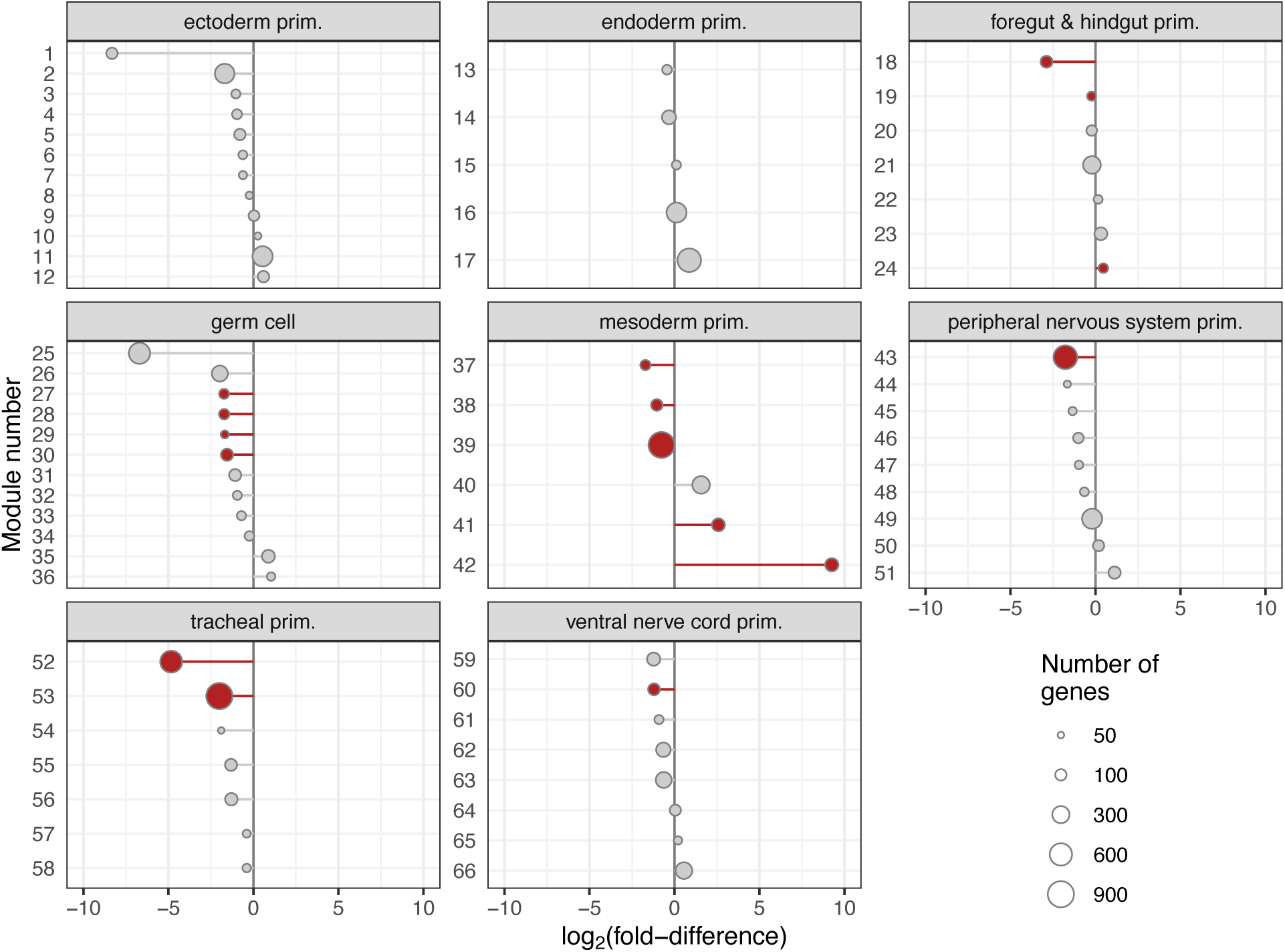
All cell-type-specific hdWGCNA modules. Points represent the module-average of the log2 fold-difference between 18°C and 25°C. Negative values indicate that the module is expressed at a higher level in 18°C-acclimated embryos and positive values indicate the module is higher in the 25°C-acclimated embryos.

**Fig. S12.**
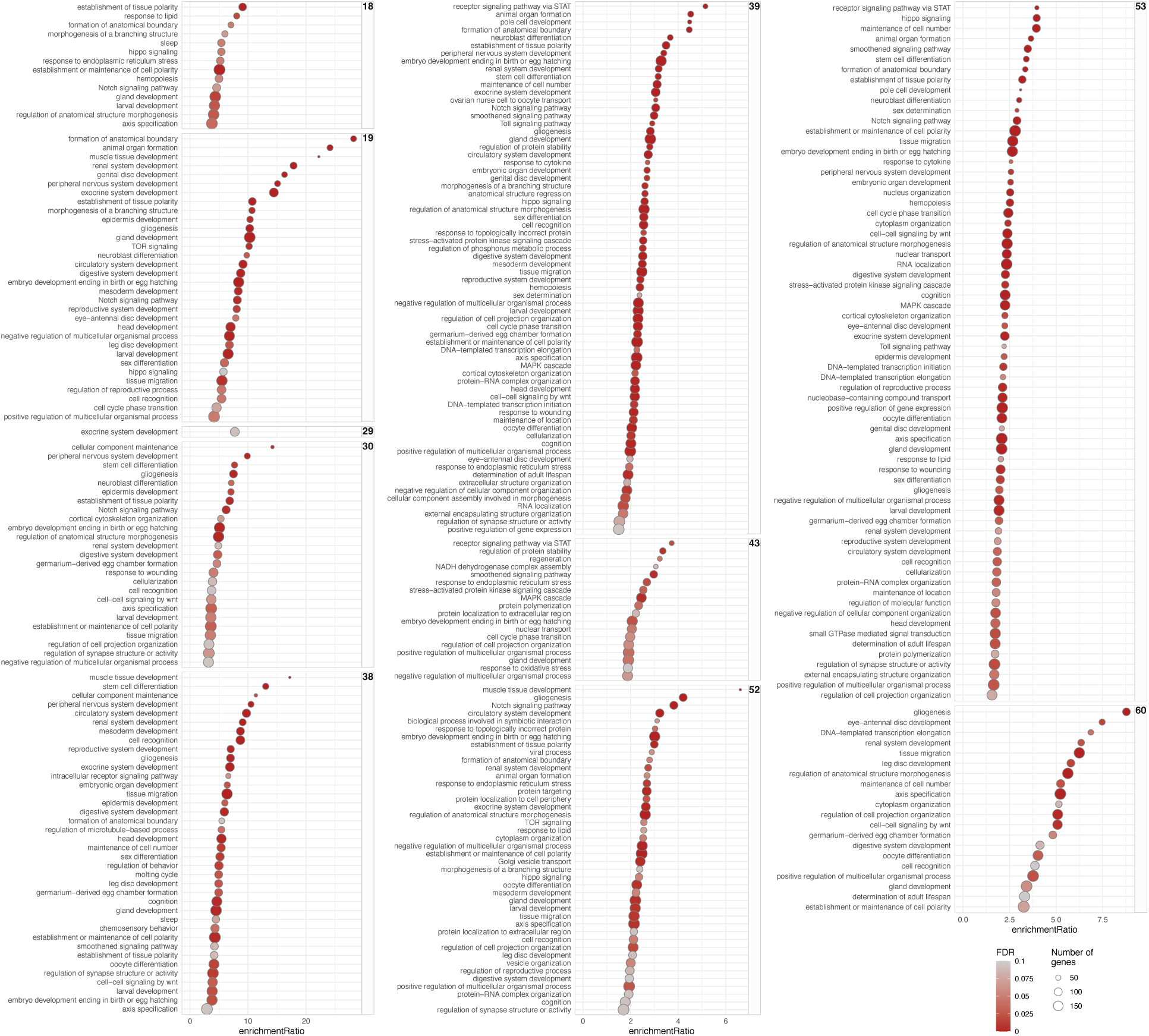
GO terms from overrepresentation analysis for all differential modules eigengenes from the cell-type-specific hdWGCNA. Points are sized by the number of genes in the GO set and colored by the FDR. Horizontal axis indicates the gene ratio (i.e., the number of DEGs present in a given GO set divided by the number of DEGs expected in that GO set by random chance, based on the background distribution of all expressed genes). All significantly overrepresented GO terms are included (FDR < 0.1).

**Fig. S13.**
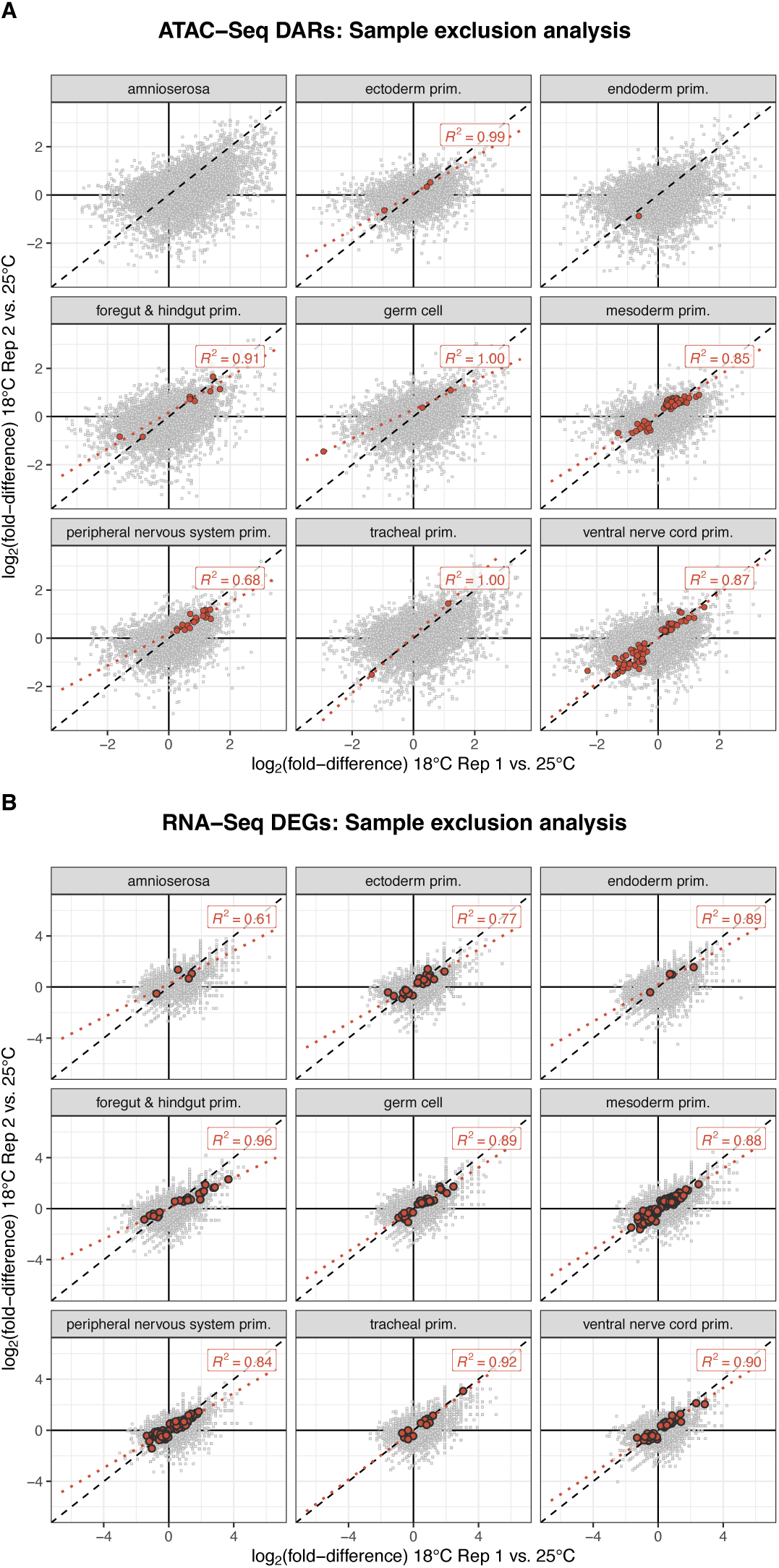
Scatter plots of the sample exclusion analysis for DARs and DEGs. Points represent individual regions or genes and their coordinate values are plotted by the log2 fold-difference in 18°C Rep 1 vs. 25°C on the horizontal axis and 18°C Rep 2 vs. 25°C on the vertical axis. Significant DARs (**A**) and DEGs (**B**) are colored in red. 1:1 line is plotted in a dashed grey line. A dotted red line represents the best fit line of the DARs and DEGs, and inset is the corresponding R^2^ value.

**Fig. S14.**
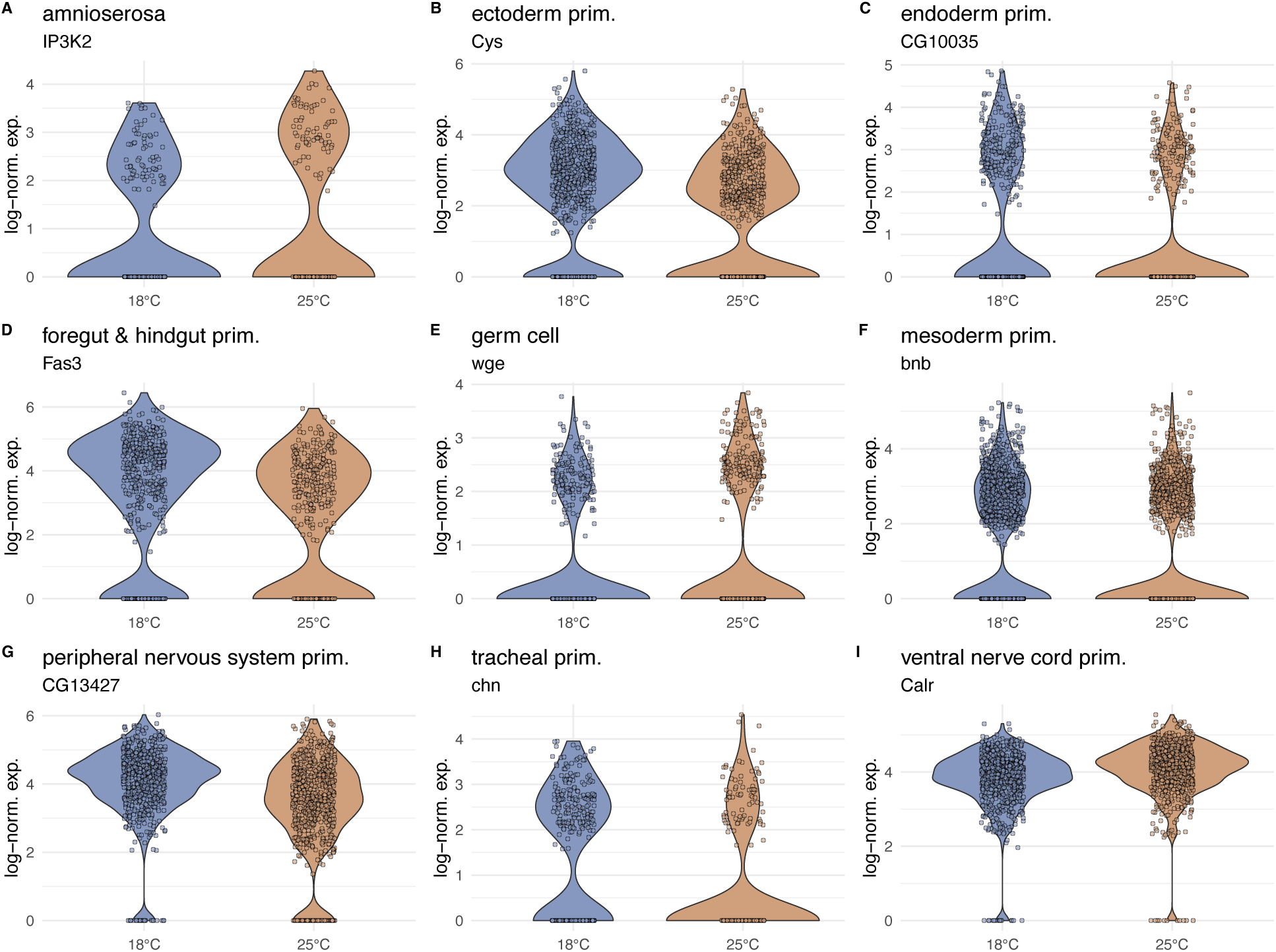
Individual violin plots of a cell-type specific DEG from each cell type. Individual dots represent the log normalized expression of the specified gene within a single nucleus.

---

**Table S1.**
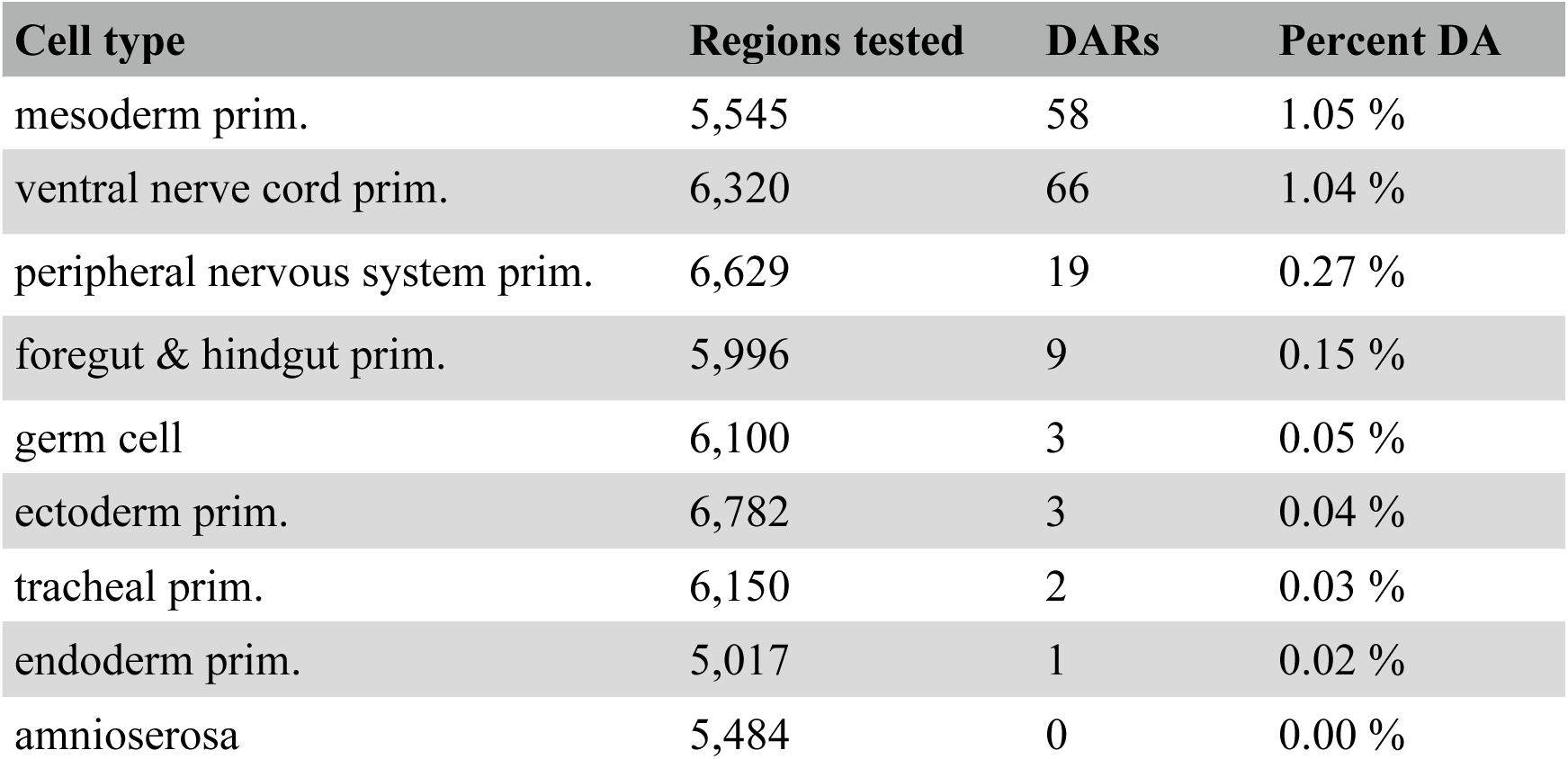
Number of regions tested for differential accessibility for each cell type (Regions tested), and number (DARs) and proportion of differentially accessible regions (Percent DA) across cell types.

| Cell type | Regions tested | DARs | Percent DA |
| --- | --- | --- | --- |
| mesoderm prim. | 5,545 | 58 | 1.05 % |
| ventral nerve cord prim. | 6,320 | 66 | 1.04 % |
| peripheral nervous system prim. | 6,629 | 19 | 0.27 % |
| foregut & hindgut prim. | 5,996 | 9 | 0.15 % |
| germ cell | 6,100 | 3 | 0.05 % |
| ectoderm prim. | 6,782 | 3 | 0.04 % |
| tracheal prim. | 6,150 | 2 | 0.03 % |
| endoderm prim. | 5,017 | 1 | 0.02 % |
| amnioserosa | 5,484 | 0 | 0.00 % |

**Table S2.** Number of genes tested for differential expression (Genes tested) for each cell type and number (DEGs) and proportion of differentially expressed genes (Percent DE) across cell types.

| Cell type | Genes tested | DEGs | Percent DE |
| --- | --- | --- | --- |
| mesoderm prim. | 1,395 | 261 | 18.71 % |
| peripheral nervous system prim. | 1,945 | 176 | 9.05 % |
| ventral nerve cord prim. | 1,651 | 60 | 3.63 % |
| germ cell | 1,711 | 34 | 1.99 % |
| foregut & hindgut prim. | 1,451 | 26 | 1.79 % |
| ectoderm prim. | 2,004 | 31 | 1.55 % |
| tracheal prim. | 1,598 | 15 | 0.94 % |
| endoderm prim. | 1,217 | 4 | 0.33 % |
| amnioserosa | 1,589 | 4 | 0.25 % |

## References

1. T. Gregor, D. W. Tank, E. F. Wieschaus, W. Bialek, Probing the limits to positional information. Cell 130, 153–164 (2007).

2. A. S. Hammonds, C. A. Bristow, W. W. Fisher, R. Weiszmann, S. Wu, V. Hartenstein, M. Kellis, B. Yu, E. Frise, S. E. Celniker, Spatial expression of transcription factors in Drosophila embryonic organ development. Genome Biol. 14, R140 (2013).

3. M. M. Harrison, A. J. Marsh, C. A. Rushlow, Setting the stage for development: the maternal-to-zygotic transition in Drosophila. Genetics 225 (2023).

4. P. Tomancak, B. P. Berman, A. Beaton, R. Weiszmann, E. Kwan, V. Hartenstein, S. E. Celniker, G. M. Rubin, Global analysis of patterns of gene expression during Drosophila embryogenesis. Genome Biol. 8, R145 (2007).

5. M. Hilker, H. Salem, N. E. Fatouros, Adaptive Plasticity of Insect Eggs in Response to Environmental Challenges. Annu. Rev. Entomol., doi: 10.1146/annurev-ento-120120-100746 (2022).

6. C. K. Mirth, T. E. Saunders, C. Amourda, Growing Up in a Changing World: Environmental Regulation of Development in Insects. Annu. Rev. Entomol. 66, 81–99 (2021).

7. O. Filina, B. Demirbas, R. Haagmans, J. S. van Zon, Temporal scaling in C. elegans larval development. Proc. Natl. Acad. Sci. U. S. A. 119, e2123110119 (2022).

8. S. G. Kuntz, M. B. Eisen, Drosophila embryogenesis scales uniformly across temperature in developmentally diverse species. PLoS Genet. 10, e1004293 (2014).

9. S. G. Kuntz, M. B. Eisen, Oxygen changes drive non-uniform scaling in Drosophila melanogaster embryogenesis. F1000Res. 4, 1102 (2015).

10. E. Gheisari, M. Aakhte, H.-A. J. Müller, Gastrulation in Drosophila melanogaster: Genetic control, cellular basis and biomechanics. Mech. Dev. 163, 103629 (2020).

11. S. Hayashi, T. Kondo, Development and Function of the Drosophila Tracheal System. Genetics 209, 367–380 (2018).

12. A. C. Martin, The Physical Mechanisms of Drosophila Gastrulation: Mesoderm and Endoderm Invagination. Genetics 214, 543–560 (2020).

13. J. Sun, F. Macabenta, Z. Akos, A. Stathopoulos, Collective Migrations of Drosophila Embryonic Trunk and Caudal Mesoderm-Derived Muscle Precursor Cells. Genetics 215, 297–322 (2020).

14. C. H. Waddington, CANALIZATION OF DEVELOPMENT AND THE INHERITANCE OF ACQUIRED CHARACTERS. Nature 150, 563–565 (1942).

15. E. Wieschaus, C. Nüsslein-Volhard, The Heidelberg screen for pattern mutants of Drosophila: A personal account. Annu. Rev. Cell Dev. Biol. 32, 1–46 (2016).

16. E. E. Mikucki, T. S. O’Leary, B. L. Lockwood, Heat tolerance, oxidative stress response tuning and robust gene activation in early-stage Drosophila melanogaster embryos. Proc. Biol. Sci. 291, 20240973 (2024).

17. B. L. Lockwood, T. Gupta, R. Scavotto, Disparate patterns of thermal adaptation between life stages in temperate vs. tropical Drosophila melanogaster. J. Evol. Biol. 31, 323–331 (2018).

18. S. P. Roberts, M. E. Feder, Changing fitness consequences of hsp70 copy number in transgenic Drosophila larvae undergoing natural thermal stress. [Preprint] (2000). 10.1046/j.1365-2435.2000.00429.x.

19. M. E. Feder, N. Blair, H. Figueras, Oviposition site selection: unresponsiveness ofDrosophilato cues of potential thermal stress. Anim. Behav. 53, 585–588 (1997).

20. J. M. Sunday, A. E. Bates, N. K. Dulvy, Global analysis of thermal tolerance and latitude in ectotherms. Proc. Biol. Sci. 278, 1823–1830 (2011).

21. B. J. Sinclair, K. E. Marshall, M. A. Sewell, D. L. Levesque, C. S. Willett, S. Slotsbo, Y. Dong, C. D. G. Harley, D. J. Marshall, B. S. Helmuth, R. B. Huey, Can we predict ectotherm responses to climate change using thermal performance curves and body temperatures? Ecol. Lett. 19, 1372–1385 (2016).

22. A. A. Hoffmann, Physiological climatic limits in Drosophila: patterns and implications. J. Exp. Biol. 213, 870–880 (2010).

23. J. H. Stillman, Acclimation capacity underlies susceptibility to climate change. Science 301, 65 (2003).

24. J. M. Sunday, A. E. Bates, M. R. Kearney, R. K. Colwell, N. K. Dulvy, J. T. Longino, R. B. Huey, Thermal-safety margins and the necessity of thermoregulatory behavior across latitude and elevation. Proc. Natl. Acad. Sci. U. S. A. 111, 5610–5615 (2014).

25. W. Tadros, H. D. Lipshitz, The maternal-to-zygotic transition: a play in two acts. Development 136, 3033–3042 (2009).

26. N. L. Vastenhouw, W. X. Cao, H. D. Lipshitz, The maternal-to-zygotic transition revisited. Development 146 (2019).

27. D. Loncar, S. J. Singer, Cell membrane formation during the cellularization of the syncytial blastoderm of Drosophila. Proc. Natl. Acad. Sci. U. S. A. 92, 2199–2203 (1995).

28. P. Tomancak, A. Beaton, R. Weiszmann, E. Kwan, S. Shu, S. E. Lewis, S. Richards, M. Ashburner, V. Hartenstein, S. E. Celniker, G. M. Rubin, Systematic determination of patterns of gene expression during Drosophila embryogenesis. Genome Biol. 3, RESEARCH0088 (2002).

29. T. Pollex, A. Rabinowitz, M. C. Gambetta, R. Marco-Ferreres, R. R. Viales, A. Jankowski, C. Schaub, E. E. M. Furlong, Enhancer-promoter interactions become more instructive in the transition from cell-fate specification to tissue differentiation. Nat. Genet., doi: 10.1038/s41588-024-01678-x (2024).

30. A. Eldar, R. Dorfman, D. Weiss, H. Ashe, B.-Z. Shilo, N. Barkai, Robustness of the BMP morphogen gradient in Drosophila embryonic patterning. Nature 419, 304–308 (2002).

31. E. M. Lucchetta, J. H. Lee, L. A. Fu, N. H. Patel, R. F. Ismagilov, Dynamics of Drosophila embryonic patterning network perturbed in space and time using microfluidics. Nature 434, 1134–1138 (2005).

32. H. F. Nijhout, M. C. Reed, Homeostasis and dynamic stability of the phenotype link robustness and plasticity. Integr. Comp. Biol. 54, 264–275 (2014).

33. J. Chong, C. Amourda, T. E. Saunders, Temporal development of Drosophila embryos is highly robust across a wide temperature range. J. R. Soc. Interface 15 (2018).

34. G. N. Somero, B. L. Lockwood, L. Tomanek, Biochemical Adaptation: Response to Environmental Challenges, from Life’s Origins to the Anthropocene (Sinauer Associates, Inc, Sunderland, MA, 2017).

35. J. Crapse, N. Pappireddi, M. Gupta, S. Y. Shvartsman, E. Wieschaus, M. Wühr, Evaluating the Arrhenius equation for developmental processes. Mol. Syst. Biol. 17, e9895 (2021).

36. M. Benet, A. Miguel, F. Carrasco, T. Li, J. Planells, P. Alepuz, V. Tordera, J. E. Pérez-Ortín, Modulation of protein synthesis and degradation maintains proteostasis during yeast growth at different temperatures. Biochim. Biophys. Acta Gene Regul. Mech. 1860, 794–802 (2017).

37. J. E. Pérez-Ortín, P. M. Alepuz, J. Moreno, Genomics and gene transcription kinetics in yeast. Trends Genet. 23, 250–257 (2007).

38. D. J. Coughlin, L. K. Nicastro, P. J. Brookes, M. A. Bradley, J. L. Shuman, E. R. Steirer, H. L. Mistry, Thermal acclimation and gene expression in rainbow smelt: Changes in the myotomal transcriptome in the cold. Comp. Biochem. Physiol. Part D Genomics Proteomics 31, 100610 (2019).

39. H. A. MacMillan, J. M. Knee, A. B. Dennis, H. Udaka, K. E. Marshall, T. J. S. Merritt, B. J. Sinclair, Cold acclimation wholly reorganizes the Drosophila melanogaster transcriptome and metabolome. Sci. Rep. 6, 28999 (2016).

40. T. N. Kristensen, H. Kjeldal, M. F. Schou, J. L. Nielsen, Proteomic data reveal a physiological basis for costs and benefits associated with thermal acclimation. J. Exp. Biol. 219, 969–976 (2016).

41. T. S. Hatakeyama, K. Kaneko, A linear reciprocal relationship between robustness and plasticity in homeostatic biological networks. PLoS One 18, e0277181 (2023).

42. H. F. Nijhout, J. A. Best, M. C. Reed, Systems biology of robustness and homeostatic mechanisms. Wiley Interdiscip. Rev. Syst. Biol. Med. 11, e1440 (2019).

43. G. N. Somero, The physiology of climate change: how potentials for acclimatization and genetic adaptation will determine “winners” and “losers.” J. Exp. Biol. 213, 912–920 (2010).

44. C. M. Sgrò, J. S. Terblanche, A. A. Hoffmann, What Can Plasticity Contribute to Insect Responses to Climate Change? Annu. Rev. Entomol. 61, 433–451 (2016).

45. D. Berrigan, Acclimation of metabolic rate in response to developmental temperature in Drosophila melanogaster. J. Therm. Biol. 22, 213–218 (1997).

46. C. J. Austin, A. J. Moehring, Local thermal adaptation detected during multiple life stages across populations of Drosophila melanogaster. J. Evol. Biol. 32, 1342–1351 (2019).

47. D. C. Hamm, M. M. Harrison, Regulatory principles governing the maternal-to-zygotic transition: insights from Drosophila melanogaster. Open Biol. 8, 180183 (2018).

48. D. Calderon, R. Blecher-Gonen, X. Huang, S. Secchia, J. Kentro, R. M. Daza, B. Martin, A. Dulja, C. Schaub, C. Trapnell, E. Larschan, K. M. O’Connor-Giles, E. E. M. Furlong, J. Shendure, The continuum of Drosophila embryonic development at single-cell resolution. Science 377, eabn5800 (2022).

49. G. Finak, A. McDavid, M. Yajima, J. Deng, V. Gersuk, A. K. Shalek, C. K. Slichter, H. W. Miller, M. J. McElrath, M. Prlic, P. S. Linsley, R. Gottardo, MAST: a flexible statistical framework for assessing transcriptional changes and characterizing heterogeneity in single-cell RNA sequencing data. Genome Biol. 16, 278 (2015).

50. E. Frise, A. S. Hammonds, S. E. Celniker, Systematic image-driven analysis of the spatial Drosophila embryonic expression landscape. Mol. Syst. Biol. 6, 345 (2010).

51. J. B. Brown, N. Boley, R. Eisman, G. E. May, M. H. Stoiber, M. O. Duff, B. W. Booth, J. Wen, S. Park, A. M. Suzuki, K. H. Wan, C. Yu, D. Zhang, J. W. Carlson, L. Cherbas, B. D. Eads, D. Miller, K. Mockaitis, J. Roberts, C. A. Davis, E. Frise, A. S. Hammonds, S. Olson, S. Shenker, D. Sturgill, A. A. Samsonova, R. Weiszmann, G. Robinson, J. Hernandez, J. Andrews, P. J. Bickel, P. Carninci, P. Cherbas, T. R. Gingeras, R. A. Hoskins, T. C. Kaufman, E. C. Lai, B. Oliver, N. Perrimon, B. R. Graveley, S. E. Celniker, Diversity and dynamics of the Drosophila transcriptome. Nature 512, 393–399 (2014).

52. H. Beati, I. Peek, P. Hordowska, M. Honemann-Capito, J. Glashauser, F. A. Renschler, P. Kakanj, A. Ramrath, M. Leptin, S. Luschnig, S. Wiesner, A. Wodarz, The adherens junction-associated LIM domain protein Smallish regulates epithelial morphogenesis. J. Cell Biol. 217, 1079–1095 (2018).

53. M. R. Johnson, R. A. Stephenson, S. Ghaemmaghami, M. A. Welte, Developmentally regulated H2Av buffering via dynamic sequestration to lipid droplets in Drosophila embryos. Elife 7 (2018).

54. Y. Barthalay, R. Hipeau-Jacquotte, S. de la Escalera, F. Jiménez, M. Piovant, Drosophila neurotactin mediates heterophilic cell adhesion. EMBO J. 9, 3603–3609 (1990).

55. M. C. Holcomb, G.-J. J. Gao, M. Servati, D. Schneider, P. K. McNeely, J. H. Thomas, J. Blawzdziewicz, Mechanical feedback and robustness of apical constrictions in Drosophila embryo ventral furrow formation. PLoS Comput. Biol. 17, e1009173 (2021).

56. J. Ray, P. R. Munn, A. Vihervaara, J. J. Lewis, A. Ozer, C. G. Danko, J. T. Lis, Chromatin conformation remains stable upon extensive transcriptional changes driven by heat shock. Proc. Natl. Acad. Sci. U. S. A. 116, 19431–19439 (2019).

57. P. K. Sorger, H. R. Pelham, Purification and characterization of a heat-shock element binding protein from yeast. EMBO J. 6, 3035–3041 (1987).

58. V. Ramalingam, X. Yu, B. D. Slaughter, J. R. Unruh, K. J. Brennan, A. Onyshchenko, J. J. Lange, M. Natarajan, M. Buck, J. Zeitlinger, Lola-I is a promoter pioneer factor that establishes de novo Pol II pausing during development. Nat. Commun. 14, 5862 (2023).

59. R. Dorfman, L. Glazer, U. Weihe, M. F. Wernet, B.-Z. Shilo, Elbow and Noc define a family of zinc finger proteins controlling morphogenesis of specific tracheal branches. Development 129, 3585–3596 (2002).

60. A. Ghabrial, S. Luschnig, M. M. Metzstein, M. A. Krasnow, Branching morphogenesis of the Drosophila tracheal system. Annu. Rev. Cell Dev. Biol. 19, 623–647 (2003).

61. C. Bravo González-Blas, S. De Winter, G. Hulselmans, N. Hecker, I. Matetovici, V. Christiaens, S. Poovathingal, J. Wouters, S. Aibar, S. Aerts, SCENIC+: single-cell multiomic inference of enhancers and gene regulatory networks. Nat. Methods, doi: 10.1038/s41592-023-01938-4 (2023).

62. M. M. Harrison, X.-Y. Li, T. Kaplan, M. R. Botchan, M. B. Eisen, Zelda binding in the early Drosophila melanogaster embryo marks regions subsequently activated at the maternal-to-zygotic transition. PLoS Genet. 7, e1002266 (2011).

63. K. N. Schulz, E. R. Bondra, A. Moshe, J. E. Villalta, J. D. Lieb, T. Kaplan, D. J. McKay, M. M. Harrison, Zelda is differentially required for chromatin accessibility, transcription factor binding, and gene expression in the early Drosophila embryo. Genome Res. 25, 1715–1726 (2015).

64. X.-Y. Li, M. M. Harrison, J. E. Villalta, T. Kaplan, M. B. Eisen, Establishment of regions of genomic activity during the Drosophila maternal to zygotic transition. Elife 3, e03737 (2014).

65. A. R. Cossins, Adaptation of biological membranes to temperature. The effect of temperature acclimation of goldfish upon the viscosity of synaptosomal membranes. Biochim. Biophys. Acta 470, 395–411 (1977).

66. M. J. Angilletta Jr, C. Condon, J. P. Youngblood, Thermal acclimation of flies from three populations of Drosophila melanogaster fails to support the seasonality hypothesis. J. Therm. Biol. 81, 25–32 (2019).

67. J. Overgaard, A. Tomcala, J. G. Sørensen, M. Holmstrup, P. H. Krogh, P. Simek, V. Kostál, Effects of acclimation temperature on thermal tolerance and membrane phospholipid composition in the fruit fly Drosophila melanogaster. J. Insect Physiol. 54, 619–629 (2008).

68. T. Enriquez, H. Colinet, Cold acclimation triggers major transcriptional changes in Drosophila suzukii. BMC Genomics 20, 413 (2019).

69. R. Morgan, J. Sundin, M. H. Finnøen, G. Dresler, M. M. Vendrell, A. Dey, K. Sarkar, F. Jutfelt, Are model organisms representative for climate change research? Testing thermal tolerance in wild and laboratory zebrafish populations. Conserv. Physiol. 7, coz036 (2019).

70. L. Comte, J. D. Olden, Evolutionary and environmental determinants of freshwater fish thermal tolerance and plasticity. Glob. Chang. Biol. 23, 728–736 (2017).

71. T. Enriquez, H. Colinet, Cold acclimation triggers lipidomic and metabolic adjustments in the spotted wing drosophila Drosophila suzukii (Matsumara). Am. J. Physiol. Regul. Integr. Comp. Physiol. 316, R751–R763 (2019).

72. D. Cheung, J. Ma, Probing the impact of temperature on molecular events in a developmental system. Sci. Rep. 5, 13124 (2015).

73. M. E. Feder, G. E. Hofmann, Heat-shock proteins, molecular chaperones, and the stress response: evolutionary and ecological physiology. Annu. Rev. Physiol. 61, 243–282 (1999).

74. J. G. Sørensen, T. N. Kristensen, V. Loeschcke, The evolutionary and ecological role of heat shock proteins. Ecol. Lett. (2003).

75. I. Belhadj Slimen, T. Najar, A. Ghram, H. Dabbebi, M. Ben Mrad, M. Abdrabbah, Reactive oxygen species, heat stress and oxidative-induced mitochondrial damage. A review. Int. J. Hyperthermia 30, 513–523 (2014).

76. X. Han, B. Sun, Q. Zhang, L. Teng, F. Zhang, Z. Liu, Metabolic regulation reduces the oxidative damage of arid lizards in response to moderate heat events. Integr. Zool., doi: 10.1111/1749-4877.12784 (2023).

77. A. Cheslock, M. K. Andersen, H. A. MacMillan, Thermal acclimation alters Na+/K+-ATPase activity in a tissue-specific manner in Drosophila melanogaster. Comp. Biochem. Physiol. A Mol. Integr. Physiol. 256, 110934 (2021).

78. J. S. Bayley, J. G. Sørensen, M. Moos, V. Koštál, J. Overgaard, Cold acclimation increases depolarization resistance and tolerance in muscle fibers from a chill-susceptible insect, *Locusta migratoria*. *American Journal of Physiology-Regulatory*, Integrative and Comparative Physiology 319, R439–R447 (2020).

79. H. A. MacMillan, G. Y. Yerushalmi, S. Jonusaite, S. P. Kelly, A. Donini, Thermal acclimation mitigates cold-induced paracellular leak from the Drosophila gut. Sci. Rep. 7, 8807 (2017).

80. G. Y. Yerushalmi, L. Misyura, H. A. MacMillan, A. Donini, Functional plasticity of the gut and the Malpighian tubules underlies cold acclimation and mitigates cold-induced hyperkalemia in Drosophila melanogaster. J. Exp. Biol. 221, jeb174904 (2018).

81. M. Rodríguez, L. Pagola, F. M. Norry, P. Ferrero, Cardiac performance in heat-stressed flies of heat-susceptible and heat-resistant Drosophila melanogaster. J. Insect Physiol. 133, 104268 (2021).

82. R. A. Krebs, M. E. Feder, Tissue-specific variation in Hsp70 expression and thermal damage in Drosophila melanogaster larvae. J. Exp. Biol. 200, 2007–2015 (1997).

83. L. B. Jørgensen, R. M. Robertson, J. Overgaard, Neural dysfunction correlates with heat coma and CTmax in Drosophila but does not set the boundaries for heat stress survival. J. Exp. Biol. 223 (2020).

84. A. L. Bryantsev, P. W. Baker, T. L. Lovato, M. S. Jaramillo, R. M. Cripps, Differential requirements for Myocyte Enhancer Factor-2 during adult myogenesis in Drosophila. Dev. Biol. 361, 191–207 (2012).

85. F. J. Blanchard, B. Collins, S. A. Cyran, D. H. Hancock, M. V. Taylor, J. Blau, The transcription factor Mef2 is required for normal circadian behavior in Drosophila. J. Neurosci. 30, 5855–5865 (2010).

86. M. Sarhan, K. Miyagawa, H. Ueda, Domain analysis of Drosophila Blimp-1 reveals the importance of its repression function and instability in determining pupation timing. Genes Cells 28, 338–347 (2023).

87. C.-Y. Nien, H.-L. Liang, S. Butcher, Y. Sun, S. Fu, T. Gocha, N. Kirov, J. R. Manak, C. Rushlow, Temporal coordination of gene networks by Zelda in the early Drosophila embryo. PLoS Genet. 7, e1002339 (2011).

88. B. Heerwaarden, V. Kellermann, C. M. Sgrò, Limited scope for plasticity to increase upper thermal limits. Funct. Ecol. 30, 1947–1956 (2016).

89. M. F. Schou, M. B. Mouridsen, J. G. Sørensen, V. Loeschcke, Linear reaction norms of thermal limits in Drosophila: predictable plasticity in cold but not in heat tolerance. Funct. Ecol. 31, 934–945 (2017).

90. S. Ramniwas, G. Kumar, D. Singh, Rejection of the beneficial acclimation hypothesis (BAH) for short term heat acclimation in Drosophila nepalensis. Genetica 148, 173–182 (2020).

91. E. Marais, S. L. Chown, Beneficial acclimation and the Bogert effect. Ecol. Lett. 11, 1027– 1036 (2008).

92. M. L. Siegal, A. Bergman, Waddington’s canalization revisited: developmental stability and evolution. Proc. Natl. Acad. Sci. U. S. A. 99, 10528–10532 (2002).

93. T. Flatt, The evolutionary genetics of canalization. Q. Rev. Biol. 80, 287–316 (2005).

94. C. K. Ghalambor, J. K. McKAY, S. P. Carroll, D. N. Reznick, Adaptive versus non-adaptive phenotypic plasticity and the potential for contemporary adaptation in new environments. Funct. Ecol. 21, 394–407 (2007).

95. C. M. Sgrò, A. A. Hoffmann, Genetic correlations, tradeoffs and environmental variation. Heredity (Edinb.) 93, 241–248 (2004).

96. S. Elena, R. Lenski, Microbial genetics: Evolution experiments with microorganisms: the dynamics and genetic bases of adaptation. Nat. Rev. Genet. 4, 457–469 (2003).

97. M. J. Angilletta Jr, T. D. Steury, M. W. Sears, Temperature, growth rate, and body size in ectotherms: fitting pieces of a life-history puzzle. Integr. Comp. Biol. 44, 498–509 (2004).

98. M. K. Burke, J. P. Dunham, P. Shahrestani, K. R. Thornton, M. R. Rose, A. D. Long, Genome-wide analysis of a long-term evolution experiment with Drosophila. Nature 467, 587–590 (2010).

99. R Core Team, R: A Language and Environment for Statistical Computing. R Foundation for Statistical Computing [Preprint] (2025). https://www.R-project.org/.

100. A. Butler, P. Hoffman, P. Smibert, E. Papalexi, R. Satija, Integrating single-cell transcriptomic data across different conditions, technologies, and species. Nat. Biotechnol. 36, 411–420 (2018).

101. T. Stuart, A. Srivastava, S. Madad, C. A. Lareau, R. Satija, Single-cell chromatin state analysis with Signac. Nat. Methods 18, 1333–1341 (2021).

102. C. Bravo González-Blas, L. Minnoye, D. Papasokrati, S. Aibar, G. Hulselmans, V. Christiaens, K. Davie, J. Wouters, S. Aerts, cisTopic: cis-regulatory topic modeling on single-cell ATAC-seq data. Nat. Methods 16, 397–400 (2019).

103. A. Thibodeau, A. Eroglu, C. S. McGinnis, N. Lawlor, D. Nehar-Belaid, R. Kursawe, R. Marches, D. N. Conrad, G. A. Kuchel, Z. J. Gartner, J. Banchereau, M. L. Stitzel, A. E. Cicek, D. Ucar, AMULET: a novel read count-based method for effective multiplet detection from single nucleus ATAC-seq data. Genome Biol. 22, 252 (2021).

104. C. S. McGinnis, L. M. Murrow, Z. J. Gartner, DoubletFinder: Doublet Detection in Single-Cell RNA Sequencing Data Using Artificial Nearest Neighbors. Cell Syst 8, 329–337.e4 (2019).

105. Y. Zhang, T. Liu, C. A. Meyer, J. Eeckhoute, D. S. Johnson, B. E. Bernstein, C. Nusbaum, R. M. Myers, M. Brown, W. Li, X. S. Liu, Model-based analysis of ChIP-Seq (MACS). Genome Biol. 9, R137 (2008).

106. C. Hafemeister, R. Satija, Normalization and variance stabilization of single-cell RNA-seq data using regularized negative binomial regression. Genome Biol. 20, 296 (2019).

107. J. A. Castro-Mondragon, R. Riudavets-Puig, I. Rauluseviciute, R. B. Lemma, L. Turchi, R. Blanc-Mathieu, J. Lucas, P. Boddie, A. Khan, N. Manosalva Pérez, O. Fornes, T. Y. Leung, A. Aguirre, F. Hammal, D. Schmelter, D. Baranasic, B. Ballester, A. Sandelin, B. Lenhard, K. Vandepoele, W. W. Wasserman, F. Parcy, A. Mathelier, JASPAR 2022: the 9th release of the open-access database of transcription factor binding profiles. Nucleic Acids Res. 50, D165–D173 (2022).

108. S. Morabito, F. Reese, N. Rahimzadeh, E. Miyoshi, V. Swarup, hdWGCNA identifies co-expression networks in high-dimensional transcriptomics data. Cell Rep Methods 3, 100498 (2023).

